# From Movement to Spread: Generating Livestock Contact Networks that Preserve Infection Dynamics

**DOI:** 10.64898/2026.09.15.751711

**Authors:** Tianjian Qin, Busra Atamer Balkan, Boris V. Schmid, Quirine A. ten Bosch

## Abstract

Animal trade links livestock holdings through contacts that change from day to day, making movement networks vital for epidemic analysis and control. Official movement records reveal these transmission routes, but privacy concerns often restrict access to the original data. Furthermore, epidemiological studies frequently require synthetic networks that accurately preserve the structural and temporal dynamics driving disease spread.

Here we introduce `NetForge`, a mechanism-informed generative framework that learns recurring sender–receiver roles from movement and farm information of Dutch national pig-movement records. We compared generators with varying structural and temporal constraints. Among them, the Operational Stochastic Block Model regime performed best by combining learned trade partner structure with constraints on how contacts persist, return, or first appear. It closely matched the accumulation of potential spreading routes in the observed network and reproduced its simulated disease transmission trajectories.

Together, we show that pairing trade structure with temporal constraints is essential for capturing infection dynamics. Because `NetForge` models trade data as a sequence of time-framed networks, it aligns well with routinely collected movement records and provides practical guidance for building more realistic synthetic movement networks from empirical trade records.

## 1 Introduction

Live-animal movements connect farms, markets, traders, and regions into temporal contact systems that can move livestock pathogens over long distances within days. Routinely collected movement records thus play an important role in veterinary epidemiology. They reveal variation in connectivity, identify animal holdings and regions that may accelerate spread, and support surveillance, tracing, and movement-control planning [1–5].

The epidemiological information in these records comes from both contact structure and contact order. Each movement records a sender, a receiver, a date, and a shipment size that can change transmission opportunity. At daily resolution, livestock trade is a temporal network in which links appear, disappear, and reappear as trading activity changes [6]. A pathogen can move along a chain only when each contact occurs after infection has reached the next sender. Contact timing, waiting periods, and recurrent trading relationships therefore determine which source–target pathways are epidemiologically feasible.

Empirical livestock studies show that temporal structure changes epidemiological interpretation. Daily cattle-trade networks in Italy displayed strong temporal organization and rapid link turnover, with consequences for invasion opportunities and surveillance priorities [7, 8]. In the United Kingdom, epidemic dynamics on a temporal cattle network differed from those inferred from static representations, even when early growth was matched [9]. Studies of German pig movements and cattle contact chains further show that time-respecting pathways can differ substantially from pathways inferred from aggregated networks [10, 11]. As a result, aggregating movements across days can introduce paths that never existed in time, compress waiting periods, and change the apparent importance of farms or regions [10, 12, 13].

These observations lead to a generative question: how can observed movement data be used to produce synthetic network panels that remain epidemiologically credible? Empirical movement records provide only one realization of a recurrent trading system. Other plausible assignments of trading partners, even when carefully constructed, may alter temporal reachability, seeding opportunities, outbreak size, and the value of surveillance targets. Contact-network uncertainty can also affect outbreak reconstruction and prediction [14, 15]. Consequently, a set of observation-driven, synthetic networks, validated for temporal structure and stress-tested for epidemic dynamics, can be very useful for preparedness analyses.

Network generators are increasingly used in livestock epidemiology to study how trading behavior shapes disease burden and to construct synthetic contact networks when empirical records are incomplete, restricted, or operationally sensitive [16–18]. A recent review emphasized that generated livestock networks should be validated for the epidemiological tasks they support, rather than by descriptive similarity alone [19]. For daily movement data, this is a demanding standard. A generator intended for an epidemic question should reproduce sparse daily activity, directed source–sink structure, repeated trading pairs, contact turnover, shipment sizes, and the time-respecting chains that determine epidemic opportunity. Related work on temporal networks likewise proposes preserving selected structural or timing features while randomizing the remaining details [20–22].

Among diverse network models, the stochastic block model (SBM) family provides a compact probabilistic language for representing recurrent livestock movement panels. It groups nodes with similar connection patterns and estimates how often contacts occur between groups. These inferred groups are called blocks. Blocks need not be densely connected communities. Instead, in a directed trade system, they can represent source, sink, intermediary, or external-pressure roles [23–25]. Layered SBM variants can infer one shared grouping across daily snapshots, while edge-valued and weighted variants can incorporate information carried by contacts [25–29]. SBMs have also been used to partition livestock trade networks and support disease-control reasoning [30].

However, applying SBMs to tackle the trade network generation problem involves two major challenges. The first challenge is that high-coverage movement records are often sparse at daily resolution, because the number of holdings is large and most do not trade on most days. Movement alone may therefore provide too little information to resolve biologically interpretable roles of the participating animal holdings. When this is the case, auxiliary information on holdings can add useful, uncertain evidence. Studies of annotated and attributed networks show that metadata can sharpen role or community inference when it is incorporated probabilistically [31–34].

A second challenge is how to carry those roles through time. Sampling each day independently from a fitted SBM can use the same daily group-level structure to generate plausible contacts while changing which trading pairs persist, reactivate, or first appear. In contrast, imposing contact-turnover constraints without a learned model of trading partners can misplace contacts in the network. A controlled comparison should examine how these sources of information affect potential disease spread.

Motivated by these challenges, we introduce `NetForge`, an SBM-guided framework for generating synthetic movement panels under observed trading conditions. We use a six-year Dutch pig-movement record collection comprising 1,792,178 transports among 9,169 farms as the foundation. One focal COROP^1^ region, Noordoost-Noord-Brabant (CR35), is represented at farm resolution, while holdings elsewhere are aggregated into regional supernodes (see Figure 1 for the rationale and Supplementary Note S2 for details). A metadata-augmented temporal SBM supplies one common grouping used to learn daily network topology, summarize the block-participation sequence and contact turnover, and model shipment sizes.

**Fig. 1.**
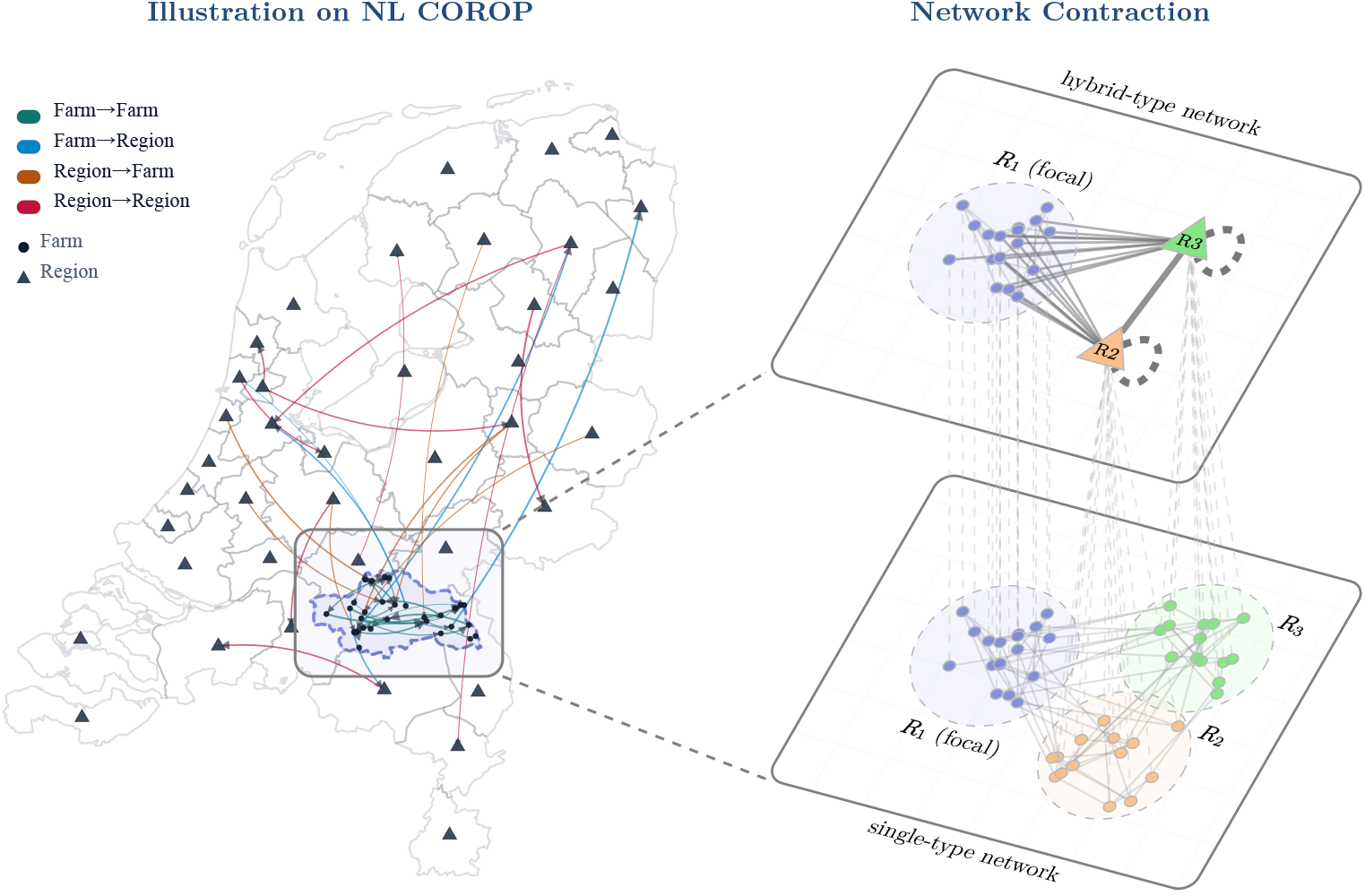
Hybrid focal-region representation. An illustration of a single-day study panel consisting of both farm nodes and regional supernodes. Edges indicate contacts. This design reduced computational cost while retaining surrounding trade flows at a coarser regional resolution. Noordoost-Noord-Brabant (CR35) was selected as the focal region because it had the largest number of recorded pig-trade events among Dutch COROP regions in the analyzed data. Farms in CR35 are kept explicit, while holdings outside CR35 are aggregated by COROP region into regional supernodes. The resulting network panel contains farm-to-farm, farm-to-region, region-to-farm, and region-to-region contact channels. Dashed self-loops around R2 and R3 indicate regional movements whose two external endpoints belonged to the same COROP region and were excluded after contraction; only movements between distinct regional supernodes were retained as region-to-region contacts.

We compare three controlled generation regimes: independent sampling of daily network snapshots from the fitted block model (*Independent SBM*); random pairing constrained by the block-participation sequence and contact turnover (*Operational Random*); and a combined generator that applies those temporal constraints to SBM-guided contacts (*Operational SBM*). After developing the workflow on the 2019 panel, we fitted and evaluated it on the 2018, 2020, and 2022 panels. We show that metadata supports a finer identification of trading roles, that plausible daily trading structure and highly correlated epidemic curves can coexist with substantial errors in temporal reachability and epidemic burden, and that the combined generator most closely reproduces both temporal pathways and simulated epidemic outcomes across the three evaluation years.

NetForge contributes both a reusable generator of synthetic network panels and an ablation framework for identifying which structural and temporal features those panels should preserve.

## 2 Results

We used `NetForge` to test whether learned trading structure and temporal constraints preserve the spreading behavior of synthetic Dutch pig-trade panels by establishing three generation regimes. We structure our analysis on the results around four core objectives: identifying whether group-level network structure reflects interpretable trading roles, assessing whether generated network panels capture plausible daily contacts, determining if these contacts form realistic time-ordered pathways, and exploring whether discrepancies in temporal structure alter simulated transmission.

Daily network statistics alone did not capture the differences in epidemic burden among generation regimes. Our analyses thus focus on connecting daily contact diagnostics to temporal reachability and transmission outcomes. Supplementary results and materials are provided in our online repository, see Supplementary Information. In addition, our companion web application, `HerdLink.nl`, enables interactive analysis of region-aggregated Dutch livestock movement networks and epidemic simulations under interventions and restrictions.

### 2.1 NetForge separates daily trading structure from temporal contact memory

We begin by introducing the framework and the logic of the regime comparison. In our framework, all three generation regimes use the same inferred block structure so that the comparison examines learned daily trading partner structure and contact-history selection within a common block representation (Figure 2). A metadata-augmented temporal SBM first learns the shared block partition from the observed movement panel and linked metadata. The joint metadata–movement inference and temporal SBM specification are described in Supplementary Note S3.

**Fig. 2.**
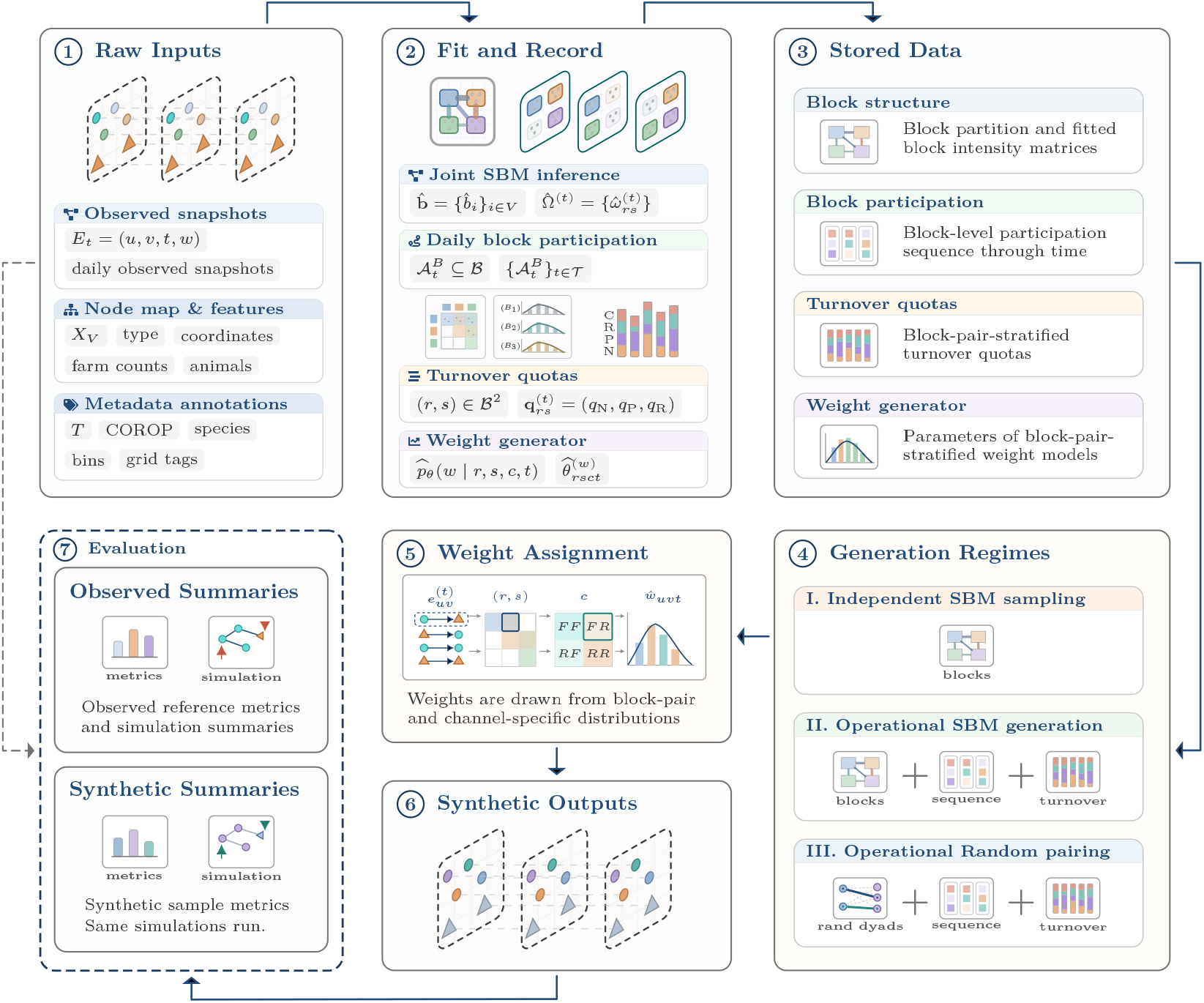
NetForge uses one learned block representation throughout generation and validation. The workflow has seven stages: (1) convert movement records into directed, weighted daily snapshots, aggregate holdings outside the focal region into regional supernodes, and annotate nodes with metadata; (2) fit a metadata-augmented SBM to the movement and annotation layers, record which blocks and their member holdings are available on each day, summarize each observed day by its active blocks, preferred number of active nodes, and contact turnover targets within each directed block pair, and fit shipment-size distributions by reusing the block pairs and hybrid channels; (3) store the data-derived information into four organized packages, all coordinated by the same inferred block structure; (4) generate panels with *Independent SBM*, *Operational SBM*, and *Operational Random*; (5) assign shipment sizes after the network topology has been selected; (6) assemble a complete node-level synthetic panel; (7) compare observed and synthetic panels using structural diagnostics and a hybrid transmission simulator.

With respect to this partition, we then derive two temporal summaries from the observed panel. The daily block-participation sequence records which inferred blocks are active on each day. Contact turnover targets record, within each directed block pair, the daily numbers of persistent, reactivated, and first-time contacts. A contact is *persistent* when it also occurred on the preceding day, *reactivated* when it occurred earlier in the panel but not on the preceding day, and *first-time* (new) when it has not appeared before. The numbers of daily active farms and regions are also stored in the turnover summary. The temporal summaries are used as eligibility rules and quotas for the operational generation regimes. Block participation defines eligible endpoint groups, contact-turnover targets guide contact selection, and active-node targets penalize shortfalls in participation counts.

The three regimes form controlled ablations. *Independent SBM* (regime I in Figure 2) samples each daily layer directly from the fitted block model, preserving learned daily contact structure without conditioning on the contact history of earlier generated days. *Operational SBM* (regime II) selects eligible contacts according to their generated contact histories from the SBM sampling pool to approach the daily block-pair-specific contact turnover targets. *Operational Random* (regime III) is constrained by these temporal summaries, but proposes sender–receiver pairs uniformly among nodes belonging to active blocks, without a learned, holding-specific daily contact structure. The three regimes compare learned trade partner structure, contact-history selection using uniform partner proposals, and their combination, thus allowing their consequences for daily network topology, temporal reachability, and transmission behavior to be assessed. The generation algorithms, contact turnover constraints, and shipment-size model are detailed in Supplementary Note S4.

### 2.2 Metadata supports a finer partition with interpretable trading roles

We next describe the trading patterns among regions and animal holdings, and examine whether metadata provide additional support for resolving their functional trading roles. Our main analysis evaluated the full workflow on a 2018 slice containing 843 farms and 38 regional supernodes across 365 daily layers. The data sources and daily panel construction methods are described in Supplementary Note S1. These 881 nodes define 775,280 possible directed non-self pairs per day. However, even the busiest day contains fewer than 500 contacts and daily all-pair occupancy remains below 0.1%. Thus, fewer than one in a thousand possible pairs is active on any day. Activity also follows a strong calendar rhythm because contact and active-node counts fall on weekends and Dutch public holidays and recover on weekdays. Consequently, the generator must place a small number of daily contacts within a large sender–receiver space while preserving recurrent activity patterns.

The focal farms are strongly connected to the surrounding trade system represented by regional supernodes. Aggregated across 2018, the panel contains 79,664 daily directed contacts on 4,329 distinct pairs, and records 24,658,750 moved pigs. Channels involving regional supernodes account for 81.0% of contacts and 90.6% of movement weight. Region–region contacts are especially concentrated where 14.2% of distinct trading pairs account for 39.7% of contacts and 59.4% of total animals moved. These results indicate that the regional network surrounding the farms is an important component of the hybrid trade system. These baseline properties are described in Supplementary Note S6, with channel-specific totals in Supplementary Table 3 and daily activity patterns in Supplementary Fig. 4.

SBM inference showed that movement edges alone supported only a coarse network division into one farm block and one regional block. In contrast, adding metadata to the contacts supported a finer partition of 11 leaf blocks at the most detailed level of the nested block hierarchy, comprising nine farm blocks and two regional blocks (see Figure 3 for the left three Sankey columns). Two higher levels of nested block structure were also identified. They progressively merged the leaf blocks into coarser interaction groups (see Figure 3 for the right two Sankey columns). Overall, metadata provided richer information that revealed otherwise hidden finer-scale block structure in the observed network panels.

**Fig. 3.**
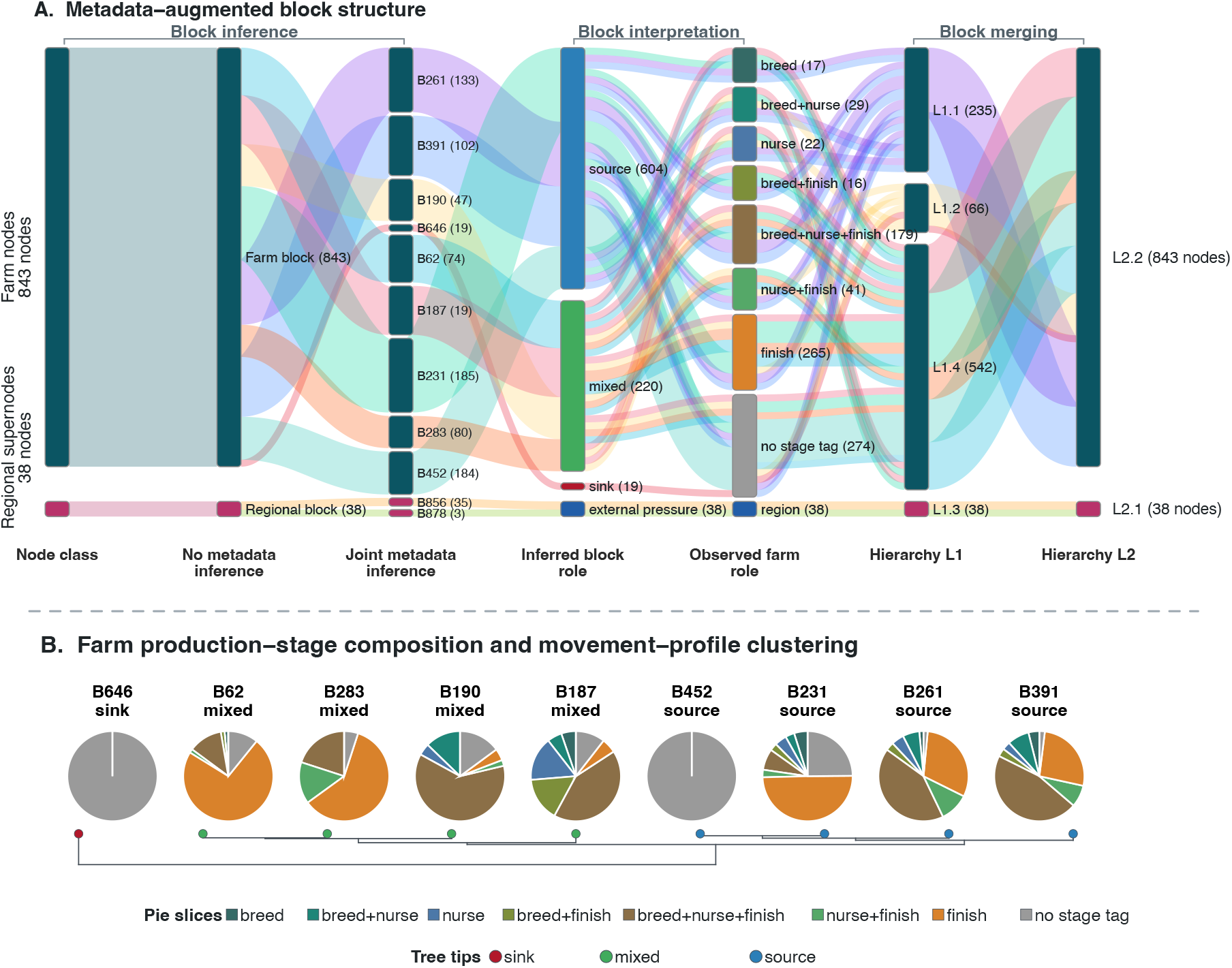
Metadata supports a finer-scale movement-role identification. **A**: The Sankey is organized into three bracketed stages. *Block inference* shows the initial node classes, the coarse no-metadata partition, and the finer joint metadata partition. *Block interpretation* connects the leaf blocks to their directed trading roles and observed farm production stages; regional supernodes are interpreted as a separate regional category because farm-stage metadata do not apply to them. *Block merging* traces the leaf blocks into hierarchy levels L1 and L2. **B**: Pies show the observed production-stage composition of the nine farm blocks, ordered by inferred role and composition. The dendrogram groups blocks by similarity in source-normalized receiving profiles, using the same distance as the corresponding column tree in Supplementary Figure 5, right panel; tip colors identify the assigned source, mixed, and sink roles.

According to metadata, the production stages of the animal holdings did not map one-to-one onto the inferred farm blocks (Figure 3-A, third and fifth Sankey columns). To interpret these inferred holding groups, we assigned each farm block a directed trading role by comparing its total outgoing and incoming shipment size. Four blocks had predominantly outgoing movement weight and were classified as sources; four had more balanced flows and were classified as mixed trading blocks; and one had predominantly incoming movement weight and was classified as a sink. The two regional blocks showed different patterns of exchange with focal farms and with other regions, indicating distinct forms of external movement pressure (Figure 3-A, fourth Sankey column).

We then compared how strongly each block connected to every other block, using both total shipment size and the proportion of each sender block’s shipment size directed to each destination. Clustering these contact profiles reproduced identical source, mixed, sink, and regional role assignments as compared to our initial interpretation (compare Figure 3-B with Supplementary Fig. 5, right panel, bottom dendrogram). Breeding and nursery farms were concentrated in several source and mixed blocks, whereas blocks containing many finisher farms included both mixed and export-oriented movement roles (Figure 3-A, middle three Sankey columns; Figure 3-B, pie charts). The role-assignment criteria, block-mixing analyses, and metadata-based interpretations are detailed in Supplementary Note S7 and Supplementary Table 4.

### 2.3 Plausible daily snapshots can still create unrealistic temporal pathways

We further examined whether the two contact proposal mechanisms, SBM sampling and uniform random pairing, placed contacts in plausible parts of the possible sender–receiver space. On a sparse example day, *Independent SBM* and *Operational SBM* reproduced the observed contact and active-node sets, whereas *Operational Random* shared no observed contacts because it sampled contact pairs uniformly without learned structure (Figure 4-A). On a high-activity day with 410 observed contacts, edge-set Jaccard similarity—the number of shared contacts divided by the size of the observed–generated union—was 0.39 for *Independent SBM* and 0.43 for *Operational SBM*, compared with 0.03 for *Operational Random* (Figure 4-B). The corresponding numbers of shared contacts were 228, 247, and 27.

**Fig. 4.**
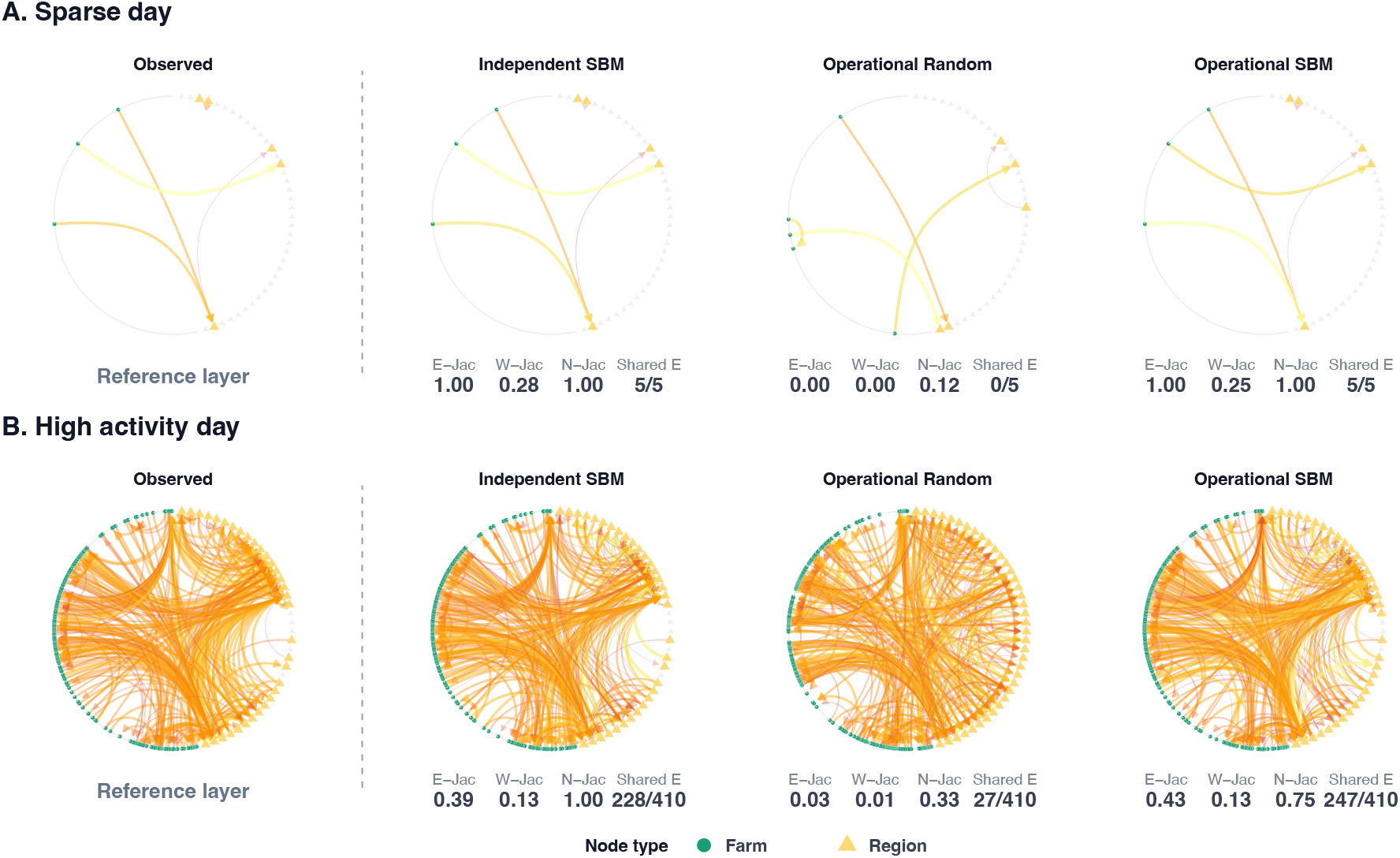
SBM-guided generation reproduces plausible daily contact structure. **A**: An extremely sparse day with five observed contacts. **B**: A high-activity day with 410 contacts. For both panels, columns compare the observed network with *Independent SBM*, *Operational Random*, and *Operational SBM* using the same node positions; circles denote farms and triangles regional supernodes. Statistics below each generated network compare it with the corresponding observed layer. E-Jac is the Jaccard overlap of directed contact pairs, W-Jac is the weighted Jaccard overlap accounting for shipment sizes, and N-Jac is the Jaccard overlap of active nodes; all range from 0 (no overlap) to 1 (complete overlap). Shared E gives the number of directed contacts present in both networks relative to the number observed that day.

Because the daily network layout is held fixed across facets in Figure 4, the high-activity day also visually shows how well each regime preserves clustered placement of contacts. Overall, SBM-guided regimes (Figure 4-B) concentrated contacts in role-structured parts of the network, closely resembling the observed pattern while still generating many plausible alternatives to the observed daily layer.

To assess how daily contacts connected through time, we introduced three complementary views (Figure 5). First, the general concept of turnover can be defined for any set observed repeatedly through time. We applied the previously defined turnover categories to directed contacts and active nodes. Contact turnover is measured on exact directed node pairs that interact; active-node turnover is measured on farms and regional supernodes that participate in trade. In each case, an element is *persistent* when it also occurred on the preceding day, *reactivated* when it occurred earlier in the panel but not on the preceding day, and *first-time* (new) when it has not appeared before. The BP (block-pair) turnover diagnostic measures contact histories within directed block pairs. Its discrepancies concern the allocation of contact histories among block pairs.

**Fig. 5.**
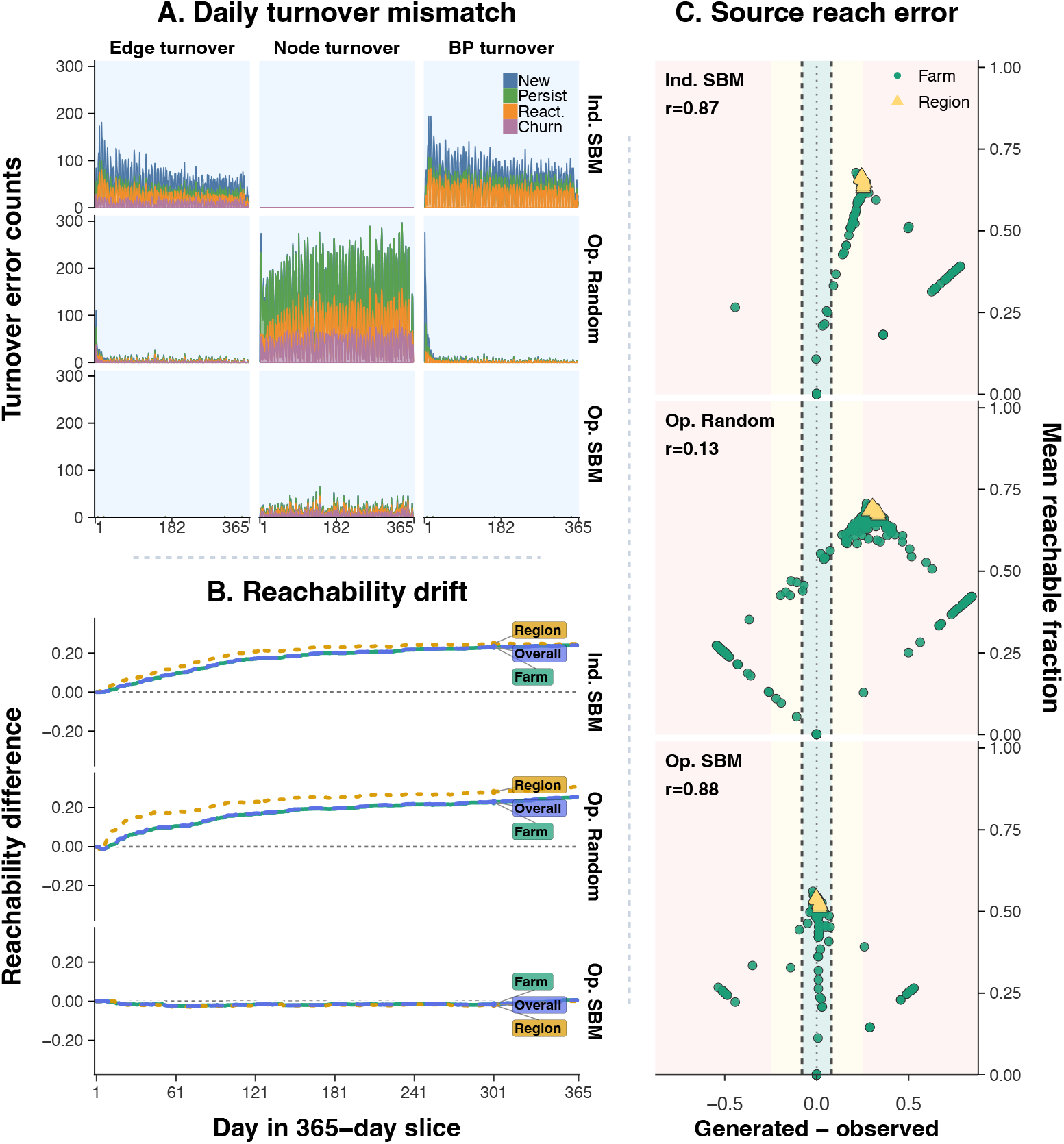
Learned contact structure and temporal memory jointly preserve temporal reachability. **A**: Daily absolute differences between generated and observed turnover counts. Columns show exact edge (contact) turnover, active-node turnover (farms and regional supernodes), and contact turnover within directed block pairs (BP turnover); rows show *Independent SBM*, *Operational Random*, and *Operational SBM*. Stacked colors separate new, persistent, reactivated, and, for edges and nodes, churn events. Churn denotes elements present on the preceding day but absent on the current day. BP turnover concerns contact histories within block pairs. Smaller counts indicate closer agreement. **B**: Temporal reachability error through the 365-day panel, with one row per generative regime. Curves distinguish farm sources, regional sources, and all sources. At each day, reachability is calculated from contacts observed up to that day as the fraction of eligible ordered source–target pairs connected by a chronologically ordered directed path. Values are generated minus observed reachability; positive values indicate excess reachable pairs and negative values indicate fewer reachable pairs. **C**: Source-level final forward-reach error under each generative regime. Each point represents a farm or regional source. The horizontal axis is generated minus observed final forward-reach fraction, while the vertical axis is their mean. The central green band denotes the closest agreement, with yellow and red bands indicating progressively larger absolute errors; dashed vertical lines mark the error thresholds. Panel annotations report the Spearman rank correlation, *r*, between observed and generated source-level forward reach.

Second, source-level forward reach is defined as the fraction of other farms and regions reachable from a given source by the end of the observation period, and in our case, after 365 days. It measures potential network-wise propagation, or downstream access, through the temporal panel.

Third, temporal reachability tracks how these pathways accumulate through time by asking how many other holdings can be reached from each source through directed contacts occurring in chronological order. The three views are further developed into visual diagnostics to identify the differences between synthetic network patterns and the observed baseline (Figure 5). Definitions and implementation details for turnover, source-level forward reach, and temporal reachability are provided in Supplementary Note S5.

*Independent SBM* preserved much of the observed ranking of sources (Spearman *r* = 0.87), but shifted many farms and regional nodes towards greater forward reach (Figure 5-C, top facet). Its generated daily layers appeared plausible; however, independent resampling changed which contacts persisted, reactivated, or first appeared, producing large contact-turnover errors overall and within block pairs, alongside excess temporal pathways (Figure 5-A, top row). Even with the same active blocks, different holding pairs can connect successive contacts into additional temporal routes. *Operational Random* reduced contact-turnover errors overall and within block pairs, but produced much larger active-node turnover errors (Figure 5-A, middle row), and its source-level reach agreement was weak (Spearman *r* = 0.13). Conditioning on the block-participation sequence and approaching contact turnover targets was therefore insufficient when candidate pairs were placed uniformly within active blocks.

*Operational SBM* gave the highest source-level agreement (Spearman *r* = 0.88) and placed most farm and regional nodes close to their observed forward reach. It also closely matched contact and block-pair turnover while keeping active-node turnover errors small. These findings suggest that temporal constraints were most informative when applied to contacts proposed from the learned block structure, but not from uniformly sampled sender–receiver pairs.

The reachability trajectories further show the cumulative consequence of these differences. *Independent SBM* and *Operational Random* both created excess temporal pathways for farm and regional sources, whereas *Operational SBM* was close to the observed reachability trajectory throughout the year. Learned daily trade partner structure and contact-history selection are therefore complementary within the preserved block-participation sequence. Omitting either component changed the ordered contacts available for spread.

### 2.4 Preserving temporal pathways recovers epidemic magnitude across yearly panels

We next asked whether discrepancies in temporal organization altered simulated transmission. We applied the same susceptible–infectious–susceptible (SIS) process to the observed panel and to panels from each generation regime (Figure 6) to compare the epidemic outcomes. The simulations served as a common stress-test of contact-panel fidelity.

**Fig. 6.**
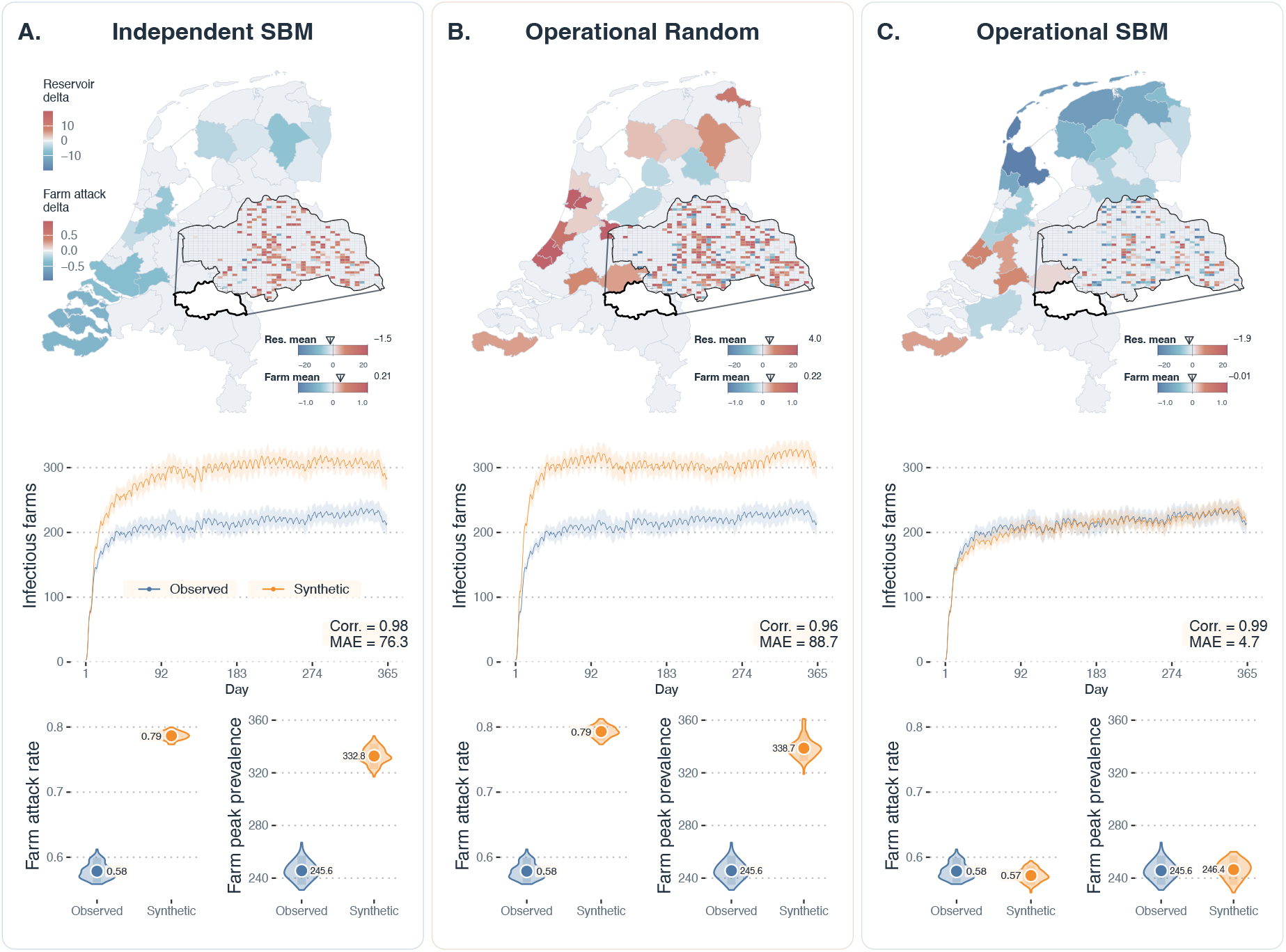
Transmission simulations distinguish the three generation regimes. Farm nodes follow an SIS process, while regional supernodes carry continuous movement-mediated pressure. Each column compares one generation regime with the observed panel using identical simulation parameters, initial seeding farms, and random-number seeds. Top-row panels show regional reservoir-pressure differences and farm-level attack-probability differences in CR35. Middle-row panels show mean daily farm prevalence with 5th–95th percentile bands. Bottom-row panels compare replicate distributions of farm attack rate and peak prevalence.

We examined three complementary aspects of performance in this stress test. Figure 6 compares the three regimes in columns A–C; within each column, the top panels assess spatial agreement in regional reservoir pressure and farm-level infection risk, the middle panel assesses the timing and magnitude of infectious-farm prevalence, and the bottom panels assess epidemic burden through farm attack rate and peak prevalence. For each focal farm, network-generated attack probability was the proportion of eligible simulations in which the farm became infected at least once, excluding simulations in which it was selected as an initial seed. Farm attack rate was the proportion of focal farms infected at least once over the full simulation horizon, counting each farm once despite possible recovery and reinfection under SIS dynamics. Peak prevalence was the largest number of simultaneously infectious focal farms within a simulation. Full outcome definitions and hybrid-simulator settings are provided in Supplementary Note S5.

*Independent SBM* and *Operational Random* produced mean prevalence curves that remained highly correlated with the observed-panel curve (Pearson *r* = 0.98 and *r* = 0.96), yet both substantially overestimated epidemic magnitude. Their prevalence mean absolute errors were 76.3 and 88.7. Both generated a mean farm attack rate of 0.79, compared with 0.58 on the observed panel, and increased mean peak prevalence to 332.8 and 338.7, compared with 245.6. These results show that a close match in curve shape can conceal substantial errors in the number of affected farms. Both regimes created excess time-ordered routes, consistent with their larger attack rates and epidemic peaks. *Independent SBM* lacked contact-history selection, whereas *Operational Random* ignored learned holding-specific trade partner choices.

In contrast, *Operational SBM* closely matched the shape and magnitude of the farm-prevalence trajectory. Its Pearson correlation with the observed mean prevalence curve was *r* = 0.99, with a mean absolute error of 4.7. Mean attack rate was 0.57, compared with 0.58 on the observed panel, and mean peak prevalence was 246.4, compared with 245.6. Mean farm-level differences in network-generated attack probability within CR35 were centered near zero. Regional reservoir pressure differences showed larger spatial biases, indicating that external contacts and regional circulation are a sensitive component of the hybrid representation.

The transmission results followed the same performance ordering of the generation regimes as the temporal diagnostics. Under the primary SIS setting with linear shipment weights, the two ablation regimes (*Independent SBM* and *Operational Random*) had excess temporal reachability and produced larger attack rates and epidemic peaks, while *Operational SBM* remained close to the observed panel in both prevalence trajectories and farm burden.

The same pattern recurred when we applied the `NetForge` workflow separately to the 2020 and 2022 validation panels (Supplementary Note S9). In both years, *Operational SBM* most closely reproduced the temporal reachability trajectory, farm attack rate, and peak prevalence. The resulting prevalence mean absolute error was 7.8 in 2020 and 6.1 in 2022, compared with 63.6 and 59.4 for *Independent SBM*, and 81.2 and 62.7 for *Operational Random*, respectively. The consistent advantage across separately fitted panels supports the repeatability of our generation strategy within the Dutch movement system.

We also tested whether this performance ordering depended on the farm-state model or on how shipment size entered the transmission hazard with the 2018 panel. We introduced two sensitivity settings that used SIS dynamics with binary contact weights and susceptible–infectious–recovered (SIR) dynamics with linear weights. *Operational SBM* was still the closest regime under both settings, although the magnitude and direction of its remaining trajectory error depended on the simulator specification (Supplementary Note S8; Supplementary Figs. 6 and 7).

## 3 Discussion

### 3.1 Trading partner choice and temporal memory across scales

Synthetic movement panels can retain plausible daily trading structure while substantially misrepresenting how many farms become infected. Our comparisons identify a combination of learned trading roles and contact-history selection that closely preserves both temporal pathways and simulated epidemic burden. This agreement emerged without directly targeting reachability or epidemic outcomes during generation and recurred across three separately fitted yearly panels. Our results link the information retained during synthesis to the epidemiological behavior of the generated networks.

`NetForge` implements this combination through block-centered network distillation. One metadata-augmented temporal SBM provides a shared representation for daily trading topology, block participation, active-node targets, contact turnover, and shipment-size models. This common representation connects constraints on trading groups to the selection of individual contacts and allows their joint consequences to be assessed.

The two ablation regimes strengthen our understanding of temporal networks. Our results show that *Independent SBM* retained much of the structure of individual trading days but changed how contacts connected across days, creating too many temporal routes. *Operational Random* approached contact-turnover targets within block pairs but paired holdings without their learned trade partner preferences, producing implausible contact placement and excess reach. Their complementary discrepancies suggest that both learned contact placement and contact-history selection are important in the movement system.

The components of `NetForge` act at different network scales. The degree-corrected SBM captures block-level organization and part of the holding-level heterogeneity in sending and receiving activity. The block-participation sequence determines which groups are eligible to trade on each day and therefore limits when group-to-group bridges can form. Within each directed block pair, contact turnover targets regulate how holding-pair contacts persist, reactivate, or first appear, adding memory to contacts between eligible groups. Different choices of these holding pairs can open or close chronological pathways even when the active blocks remain the same. Contact-history selection combined with learned partner structure thus brings *Operational SBM* closer to the observed network across scales. The performance of *Operational Random* also shows that holdings within a block are not fully interchangeable, because uniform pairing ignored holding-specific sending and receiving propensities even though the generation regime used the observed block-participation sequence and contact turnover targets.

The persistent, reactivated, and first-time categories provide a compact description of holding-pair contact memory, as described in Results Section 2.3 and detailed in Supplementary Notes S4 and S5. This connects our approach with dynamic-network models that distinguish relationship formation from persistence, and with studies showing that network memory can operate over several time scales [35–37]. Repeated contacts within a group may also reflect stable partner preferences, which is more than temporal memory alone [38].

In our present study, *Operational SBM* combined both learned block structure and inherited temporal memories. Looking ahead, a next extension would be to model the duration of inactive periods and the recurrence of trading partnerships directly, allowing block-participation and contact turnover to be generated instead of inherited from an observed panel.

### 3.2 A conditional generator of movement networks

`NetForge` addresses a different setting from generators designed for systems with little or no movement data. For example, maximum-entropy approaches estimate swine movements from operation type, size, and distance, while mechanistic models represent trading decisions and examine how changes in trade affect disease dynamics and control [16–18, 39]. By contrast, `NetForge` generates synthetic contact realizations conditioned on an observed block-participation sequence, as described in Results Section 2.1 and Supplementary Note S4.

Our study benefits from an unusually information-rich dataset containing six years of national movement records that were linked to farm locations and agricultural-registry metadata [40, 41]. This creates an opportunity to further quantify the methodological robustness under incomplete observation scenarios. For example, sensitivity studies could mask movements or metadata under random missingness and systematic patterns tied to holding type, region, or time period. Refitting the full workflow to the remaining records and comparing it with the unmasked panel would quantify how incompleteness affects inferred trading roles, generated contacts, temporal reachability, and transmission outcomes. It would also test how much missing structure can be reconstructed when the holding population and part of its metadata are still known [19].

*Operational SBM* conditions on the observed block-participation sequence and on block-pair-specific contact-turnover targets for each day while allowing the realized holding pairs to vary. It is therefore suited to examining contact uncertainty under a known participation pattern. Reconstructing missing contacts among known holdings is an appealing and feasible next step. However, extrapolating to entirely unobserved holdings, regions, or periods would require additional models for participation, activity, and turnover. Predicting changes in trading activity in another year, after a market disruption, or under movement restrictions thus lies beyond the present study. Related temporal-network generators similarly differ in whether they reproduce selected temporal patterns or learn the processes that generate those patterns [21, 22, 42].

### 3.3 Temporal pathways and infection dynamics

Temporal reachability provided a useful connection between structural and epidemiological validation. It showed how time-ordered routes accumulated from daily contacts and separated the two ablation regimes from *Operational SBM* before disease-specific assumptions were introduced (Results Section 2.3; Supplementary Note S5). This use of temporal path structure is consistent with work showing that first-arrival distances and related temporal-path measures can distinguish networks with different spreading properties [43].

However, reachability is only a screening measure that records whether a chronological route exists but not shipment size, transmission probability, or the finite period during which an infected holding can transmit. Consequently, transmission simulations are still needed to determine whether the temporal-structural differences alter epidemic outcomes under more sophisticated settings and mechanisms.

Our main SIS analysis adds an important validation lesson (Results Section 2.4; Supplementary Note S5). *Independent SBM* and *Operational Random* both retained a similar prevalence-curve shape because they matched much of the daily activity pattern. However, their excess temporal pathways produced larger attack rates and epidemic peaks. Curve correlation alone can therefore conceal important differences in prevalence magnitude and outbreak burden. A robust validation should compare trajectory magnitude, scalar outcomes, replicate variation, and spatial allocation alongside temporal correlation.

The sensitivity analyses retained *Operational SBM* as the closest regime while clarifying how the remaining trajectory gaps depended on the downstream model (Supplementary Note S8). SIR dynamics with linear shipment weights reduced absolute trajectory differences for all three regimes, with *Operational SBM* showing a small early underestimation. Binary weighting largely reduced the larger gaps of the two ablation regimes but modestly increased the remaining *Operational SBM* trajectory gap; *Operational SBM* nevertheless remained closest in simulated trajectory and epidemic burden. We conclude that shipment-size differences contributed to the quantitative bias of the ablations, while contact placement and timing remained important when every movement was given equal weight.

### 3.4 Data sharing and disclosure risk

A major motivation for generating synthetic movement networks is to enable data sharing and analysis when the original records are sensitive. However, the exact features that make synthetic data useful for epidemiology–such as specific trading roles, repeated partnerships, and temporal patterns–can also act as unique signatures for individual holdings. As seen in human interaction networks, distinctive local structures and temporal habits can sometimes allow attackers to re-identify nodes even in anonymized datasets [44, 45].

This creates a natural tension between analytical utility and privacy. In a supplementary benchmark for the 2018 CR35 panel, we tested whether pig-stock ranks could identify anonymous farm nodes through their ranks in annual contact counts, total shipment volume, or distinct incoming and outgoing partner counts. *Operational SBM* combined close epidemiological agreement with exact identity-linkage precision of 0.119–0.429% across these measures, using either known network membership or the full registry and network rosters (Supplementary Note S10; Supplementary Table 5). Thus, fewer than one in 200 attempted identity assignments were expected to be correct, even though substantial stock–activity associations were retained. Relatively high associations did not make individual holdings readily identifiable by the rank-matching approach.

Nevertheless, our results do not yet guarantee full anonymity, and any public release of synthetic networks should carefully weigh disclosure risks. Future studies could evaluate more sophisticated attacks, particularly those that combine external registry attributes with local trading structures and temporal profiles [45]. It is also crucial to test whether sensitive attributes can be inferred or if accessing multiple synthetic panels increases risk [46]. Ultimately, our block-centered approach offers a foundation for integrating formal privacy controls on network structure, activity, and shipment summaries [47]. Testing these in-depth controls against both privacy metrics and epidemiological outcomes will be an important next step.

### 3.5 A staged validation strategy

Our results support a staged approach to validating synthetic temporal networks. Daily diagnostics assess whether individual snapshots are plausible. Turnover and temporal reachability show how those contacts accumulate into chronological pathways. Transmission simulations then stress-test whether the resulting networks preserve epidemic dynamics. A disclosure assessment addresses the separate question of what information a proposed release may reveal.

Across three yearly movement panels, learned partner structure and contact-history selection together closely preserved spreading behavior that either component alone failed to reproduce. `NetForge` connects these ingredients through one shared block representation, providing a practical basis for studying contact uncertainty under observed trading conditions. A methodological lesson is that network synthesis should be evaluated by the pathways and downstream processes that the generated contacts support, as well as by the appearance of individual snapshots.

## 4 Methods

### 4.1 Data preprocessing

#### 4.1.1 Movement data and daily panels

We used anonymized Dutch Identification and Registration pig-movement records covering six calendar years (2018–2023), together with farm-location and agricultural-registry metadata. Each movement was mapped to a sender, receiver, calendar day, and shipment size. Records with identical ordered endpoints on the same day were combined into one weighted contact by summing shipment sizes. The system was treated as a sequence of directed, weighted daily graphs. Movement order within a day was therefore not modeled. The complete 2019 panel was used to develop the workflow and fix all analysis choices. Data-derived components were subsequently refitted separately to the complete 2018, 2020, and 2022 panels. In our study, the 2018 panel served as the primary evaluation, while the same prespecified main analyses were applied to the 2020 and 2022 panels as additional temporal validation tests. These additional results are reported in Supplementary Note S9. The 2021 and 2023 panels each lacked one snapshot within a full year and were excluded due to incomplete daily coverage. Data sources, aggregation rules, and calendar definitions are described in Supplementary Note S1.

#### 4.1.2 Hybrid farm–region representation

We used a hybrid network representation of the Dutch pig-movement system comprising two types of nodes. We chose the COROP region CR35 (Noordoost-Noord-Brabant) as the focal region and kept every animal holding in CR35 as an explicit farm node. Animal holdings outside CR35 were aggregated by COROP region into regional supernodes, and movements among external regions were represented as region-to-region contacts. This produced four directed channels: farm-to-farm, farm-to-region, region-to-farm, and region-to-region. The representation preserves focal farms for farm-level outcomes while retaining external import, export, and circulation at regional resolution. CR35 was selected because it ranked highest among Dutch COROP regions in both the number of pig farms and the number of recorded trade events in the analyzed data. Supplementary Note S2 describes the contraction and node attributes.

### 4.2 Modeling approach and temporal summaries

#### 4.2.1 Metadata-augmented temporal stochastic block model

In our framework, we first fitted a directed, degree-corrected, nested temporal SBM to the daily movement layers using the inference routines implemented in `graph-tool` [24, 25]. An additional annotation layer linked farm and regional supernodes to categorical or discretized metadata tags. The annotation tags used in our study were derived from the Geographical Information System of Agricultural Businesses (GIAB) dataset [40, 41] that described region, coordinate source, production stage, farm type, animal and farm counts, and coarse spatial location. We imposed node-class constraints that kept farms, regional supernodes, and metadata tags in separate leaf blocks during block-structure inference, while allowing their branches to merge at higher levels of the hierarchy.

Inference selected the nested partition with the shortest description length and produced one leaf-block assignment for the data nodes. Metadata edges contributed to this assignment but were excluded from generated trade panels, movement diagnostics, and transmission routes. The SBM modeled whether contacts occurred; shipment sizes were fitted after the partition had been inferred. Full graph construction and fitting details are provided in Supplementary Note S3.

#### 4.2.2 Block-centered network distillation

After inference, we summarized the observed panel in terms of the leaf blocks. For each day, we stored the active block set, preferred numbers of active farm and regional nodes, daily contact totals, and the observed counts of persistent, reactivated, and first-time (new) contacts in every directed block pair. A contact was persistent if it also occurred on the previous day, reactivated if it occurred earlier in the panel but not on the previous day, and first-time if it had not occurred earlier. Churn was the set of previous-day contacts absent on the current day. Shipment sizes were summarized after topology selection using hierarchical count models fitted separately for each hybrid channel. Parameters were estimated from specific day–block–pair cells and progressively pooled toward channel, day, and global levels. Within each hybrid channel, estimates for a specific day and directed block pair were partially pooled with estimates from larger data groups. This prevented one or two unusual shipments from dominating sparse cells, while cells with many observations kept their characteristic shipment-size patterns. When an exact combination was absent, the most specific available pooled estimate was used. The reported shipment-size models used a shifted negative-binomial family for positive integer shipment sizes. Overall, reusing one partition for observed and generated panels made block-pair targets directly comparable and aligned the network topology, activity, turnover, and shipment-size modules.

### 4.3 Synthetic network generation

After fitting the SBM and deriving the temporal summaries, we generated synthetic network panels under three alternative regimes. These regimes form parallel branches of the workflow. All three reuse the fitted block partition, but differ in their trade partner choices and contact-history selection.

#### 4.3.1 Independent SBM regime

*Independent SBM* sampled each daily layer from the corresponding fitted SBM without conditioning on earlier generated days. It preserved the daily block structure encoded by the model but imposed no explicit constraints on contact persistence, reactivation, or first appearance across days. The observed daily block-participation sequence was not enforced but implicitly realized by daily SBM sampling, without a separate activity filter.

#### 4.3.2 Operational Random and Operational SBM regimes

The other two branches applied explicit block-eligibility rules, used observed active-node counts to penalize participation shortfalls, and approached persistent, reactivated, and first-time contact targets within each directed block pair. *Operational Random* sampled candidate pairs uniformly among nodes in active blocks. It did not draw its proposals from daily SBM layers. *Operational SBM* instead accumulated candidates through repeated draws from the corresponding fitted SBM layers.

In both regimes, candidates with inactive endpoint blocks were removed. The remaining candidates were classified according to their contact history in the synthetic panel, and selected within directed block pairs to approach the turnover targets. Scarce block-pair and turnover categories were processed first. Proposal support and reductions in active-node count shortfalls guided selection; counts above the targets incurred no penalty. Unmet quotas were recorded, and remaining candidate capacity was used to approach the daily contact total. Supplementary Note S4 gives the full selection sequence.

### 4.4 Assessment through movement and spread

#### 4.4.1 Daily and cross-day network validation

Observed and generated network panels were compared at daily, block-pair, node, and whole-panel scales. The diagnostic set included contact and active-node counts, hybrid-channel shares, degree and strength distributions, contact turnover, active-node turnover, contact turnover within directed block pairs, and contact and node overlap. Temporal reachability was calculated across the ordered daily layers. The idea is that a source was considered to reach a target when a directed path respected contact order, traversed at most one contact per day, and could wait across days. We compared source-level final forward reach for each source and the daily accumulation of reachable ordered pairs for farm, regional, and all sources. These temporal measures detect temporal pathways that are not visible in marginal snapshot summaries. Definitions and additional diagnostics are in Supplementary Note S5. A supplementary comparison of GIAB pig-stock rank with holding-level activity ranks, together with a rank-based identity-linkage benchmark, is described in Supplementary Note S10.

#### 4.4.2 Simulation-based transmission behavior assessment

A stochastic hybrid transmission simulator was applied to the observed panel and each synthetic panel. Regional supernodes carried nonnegative movement-mediated pressure, while focal farms followed compartmental infection dynamics. Farm-to-farm contacts transmitted directly between holdings, farm-to-region contacts seeded regional pressure, region-to-farm contacts exposed susceptible farms, and region-to-region contacts propagated pressure to receiving regional reservoirs.

Our main comparison used susceptible–infectious–susceptible (SIS) farm dynamics with linear shipment weighting. Under SIS dynamics, infectious farms recovered to the susceptible state and could therefore become infected again. Shipment sizes entered the contact hazards linearly after division by a positive scale estimated from the observed panel and then held fixed across observed and generated panels.

We evaluated two additional sensitivity settings based on the main setting. The first replaced SIS dynamics with susceptible–infectious–removed (SIR) dynamics, under which recovered farms could not be reinfected. The second replaced linear shipment weighting with binary contact weights, so that every recorded movement contributed unit contact intensity regardless of shipment size. All remaining epidemiological parameters, generated panels, initial farm seed sets, random-number seeds, and replicate settings were held fixed across the three simulation settings.

We compared daily infectious-farm prevalence and its replicate intervals, farm attack rate, peak prevalence, farm-level network-generated attack probability, and regional reservoir pressure in our main results. Each generation regime was represented by 10 independently generated panels, each evaluated with 10 matched epidemic replicates; the observed panel was evaluated with the same 10 initial-seed sets and random-number seeds. Additional results are reported in our online repository, see Supplementary Information. The simulator was used as a common stress test of contact-panel fidelity without being calibrated to pathogen-specific parameters. The simulator equations, outcome definitions, and sensitivity settings are described in Supplementary Note S5, and the sensitivity results are reported in Supplementary Note S8.

## Data availability

The holding-level movement records and metadata contain sensitive information and cannot be released publicly under the data provider’s access conditions. Derived summary tables used in the figures, simulation outputs, and representative synthetic network realizations that pass disclosure review will be released through our online repository, see Supplementary Information.

## Code availability

The implementation and the scripts used for network generation, diagnostics, transmission simulation, analyses, and visualization are available in our online repository, see Supplementary Information. A permanent repository URL and archived software release will be provided with the accepted article.

## Acknowledgements

We thank the Netherlands Enterprise Agency (RVO) and the custodians of the agricultural census and GIAB registry for providing access to the movement and farm metadata.

## Supplementary Information

The Supplementary Information provides detailed data construction, model definitions, generation algorithms, and additional results. It is available at https://git.wur.nl/synthetic-movement-network/paper. The codebase is available at https://git.wur.nl/synthetic-movement-network/NetForge.

## Footnotes

1 A COROP region is a statistical region of the Netherlands.

