## Supplementary for "From Movement to Spread: Generating Livestock Contact Networks that Preserve Infection Dynamics"

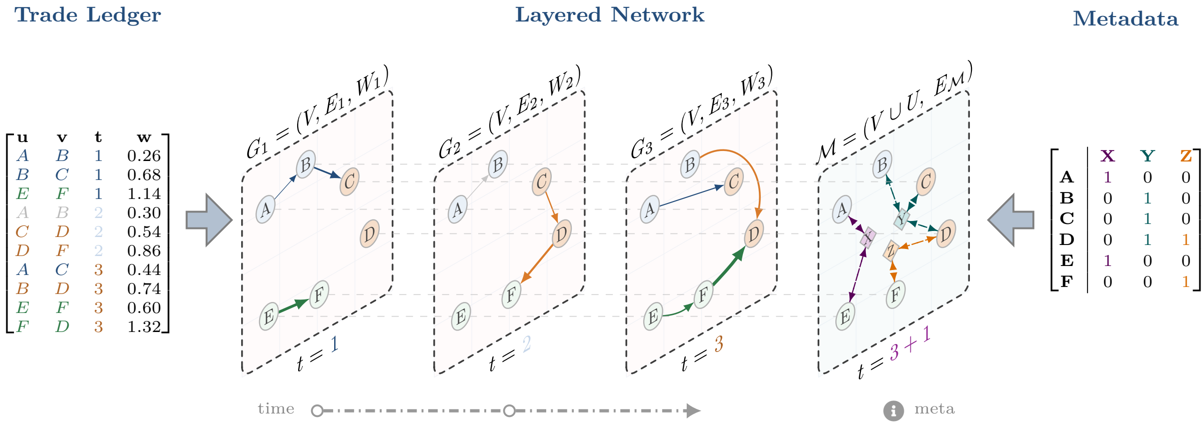


**Supplementary Fig. 1. Construction of daily movement and metadata layers.** Movement records are aggregated into directed, weighted daily contacts, while farm attributes are encoded as annotation links used during joint block inference. Node labels, edge weights, and metadata fields are illustrative.

### Supplementary Methods

#### Supplementary Note S1: Daily movement panel and metadata construction

We reconstructed pig movement networks from two Dutch national data sources. Movement events were derived from the Dutch Identification and Registration system maintained by the Netherlands Enterprise Agency (RVO), and farm-location and farm-attribute information was obtained from the Dutch agricultural census and GIAB geospatial registry [40, 41]. The full reconstructed movement records covered six years of dated, directed movements among pig holdings. The 2019 slice was used to develop the workflow and fix the analysis choices. For the reported 2018 analysis, the SBM, block-participation sequence, contact turnover targets, and shipment-size model were fitted again to the 2018 panel. The same procedures were applied also to the 2020 and 2022 panels.

Each movement record was mapped to a source node, target node, calendar day, and shipment size. Multiple records with the same ordered endpoints on the same day were aggregated by summing shipment sizes, so the daily edge list contains one edge $(u, v, t, w_{uvt})$ for each realized directed contact on day $t$. The weight $w_{uvt}$ is the total shipment size on that directed movement day. Calendar time was represented as an ordered sequence of daily layers, $\mathcal{T} = \{t_1, \ldots, t_L\}$, where $L = 365$ for 2018 and 2022, and $L = 366$ for 2020. The daily graph is

$$G_t = (V, E_t, W_t),$$

where $V$ is the node universe, $E_t \subseteq V \times V$ is the directed edge set on day $t$, and $W_t = \{w_{uvt} : (u, v) \in E_t\}$ contains positive movement weights. Thus $G_t$ is a discrete-time dynamic graph. Supplementary Fig. 1 illustrates the snapshot construction and the separate metadata layer.

#### Supplementary Note S2: Hybrid farm–region representation

We used a hybrid farm–region representation to model focal-region dynamics under external movement pressure. Let $F$ denote explicit farms inside the focal COROP region and let $R$ denote external COROP supernodes. All holdings inside the focal region remain individual farm nodes. Holdings outside the focal region are contracted by COROP region into regional supernodes. The node set is therefore

$$V = F \cup R,$$

with node type $\tau(v) \in \{F, R\}$. A movement between two focal farms is represented as a farm-to-farm edge. A movement from a focal farm to an external holding becomes a farm-to-region edge. A movement from an external holding to a focal farm becomes a region-to-farm edge. A movement between two external holdings becomes a region-to-region edge when both endpoints are represented by external supernodes. These four ordered channels are denoted

$$\mathcal{K} = \{F \to F, F \to R, R \to F, R \to R\}.$$

For each edge $(u, v) \in E_t$, the channel is $k(u, v) = (\tau(u), \tau(v))$.

This representation is asymmetric in resolution by design. It retains focal farms as explicit premises for farm-level analysis and compresses external holdings into regional import, export, and circulation reservoirs. The regional supernodes aggregate movement interfaces between the focal region and the rest of the Dutch pig movement system.

The choice was motivated by two considerations. First, full national farm-level inference and repeated posterior generation are computationally expensive because candidate dyad sets, turnover pools, statistical evaluations, and transmission simulations all scale rapidly as the explicit node set expands. Second, data governance constraints often make a coarser external representation more realistic for applied focal-region analyses in which investigators have detailed local data but only aggregated or estimated information about surrounding regions.

In this study, we chose CR35 (Noordoost-Noord-Brabant) as the focal region and retained all farms inside it. CR35 has the highest numbers of pig farms and recorded trade events among the Dutch COROP regions in the analyzed data. Fig. 1 illustrates the contraction from a full farm-level network to the hybrid representation. Movements whose two external endpoints belonged to the same COROP region were excluded after contraction, and only movements between distinct regional supernodes were kept as region-to-region contacts.

#### Supplementary Note S3: Metadata-augmented temporal SBM

Once the daily hybrid panel and metadata layer have been constructed, we fit a temporal SBM to infer groups of nodes with similar movement positions. Each data node is assigned to a latent block through unsupervised inference. Nodes in the same block may send to similar targets, receive from similar origins, or interact with regional supernodes in similar ways. The model is layered because it uses one movement layer per day, directed because sender and receiver roles differ, degree-corrected because activity varies within blocks, and nested because detailed groups can merge into broader groups. We use the most detailed, or leaf, blocks as inferred movement roles.

The fitted graph contains daily movement edges and a separate set of metadata annotation edges. Metadata tags represent coarse attributes, for example, COROP region, coordinate source, farm-type and production descriptors, binned farm and animal counts, and coarse location cells. In the directed fit, annotation links are added in both directions and assigned to their own layer. They provide node-side evidence for grouping rarely active holdings but are excluded from generated trade edges, structural movement diagnostics, and transmission routes.

Movement structure and metadata support are fitted jointly (Supplementary Fig. 2). Farm nodes and regional supernodes cannot share a leaf block, although they can merge at higher levels of the hierarchy. Leaf farm blocks describe heterogeneity among explicit CR35 farms and leaf regional blocks describe external movement roles.

We fit the nested layered SBM with the nonparametric inference routines in `graph-tool` [24, 25]. The topology fit models whether directed contacts occur. Shipment sizes are fitted after the partition is inferred, and the resulting node-to-block assignment is reused for all downstream generation components.

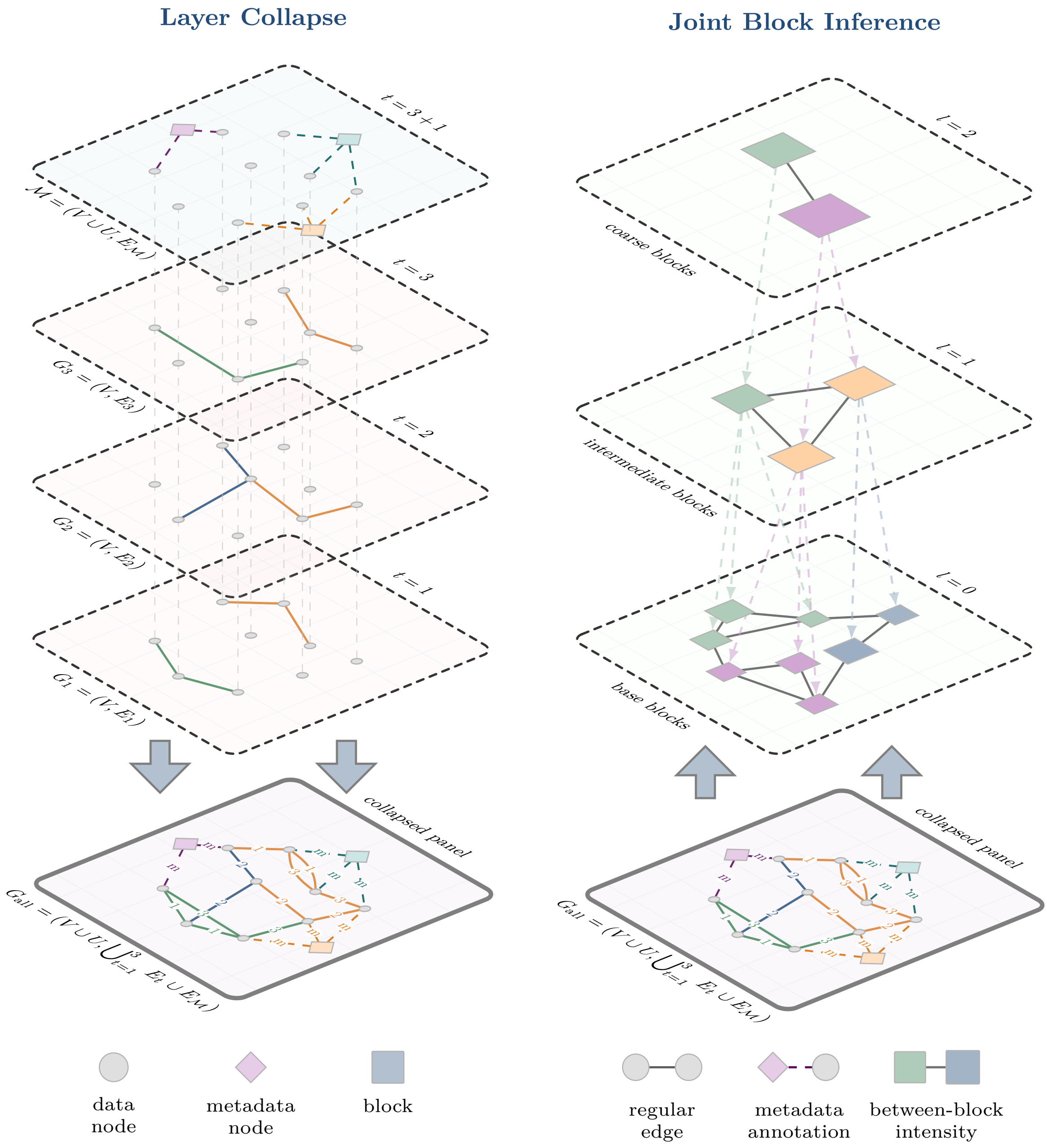


**Supplementary Fig. 2. Joint inference from temporal movement layers and metadata annotations.** Movement layers and the metadata layer share the same data nodes; metadata tags contribute node-side evidence but are excluded from the generated movement panel.

#### Supplementary Note S4: Shared-block temporal generation and shipment sizes

After fitting, we summarize the observed panel using the inferred groups. This block-centered network distillation records four quantities for each day: the directed contact pattern learned by the SBM, the groups that are active and the preferred number of active nodes, the numbers of persistent, reactivated, and first-time contacts within each directed group pair, and shipment-size distributions indexed by day, group pair, and hybrid channel.

The same node groups are used on both sides of the comparison. They are calculated from the observed network, then used to classify candidate contacts and assign weights in every generated network. This common grouping stabilizes estimates for sparse node pairs and ensures that a source or receiver group has the same meaning in the topology, activity, turnover, and shipment-size components.

We implement three generation regimes throughout the study. *Independent SBM* samples each daily layer from the fitted layered SBM without an enforced observed block-participation sequence or contact-history selection. Its realized block-participation sequence is nevertheless preserved. We use *operational* for the two regimes that assemble each daily layer through explicit activity and turnover rules. *Operational Random* proposes eligible node pairs uniformly, whereas *Operational SBM* samples candidate contacts from the fitted SBM layers.

The central operation in the two operational regimes is to reproduce the observed mix of contact histories within each directed group pair on each day. A contact is *persistent* when it also occurred on the previous day, *reactivated* when it occurred earlier in the panel but not on the previous day, and *new* when it has not appeared earlier. Churn is the set of contacts present on the previous day but absent on the current day; its count follows from the selected persistent contacts. These categories and the associated time-respecting paths are illustrated in Supplementary Fig. 3.

For each observed day, the operational generator also stores which groups are active and the preferred number of active farm and region nodes. A proposed contact is eligible only when both endpoint groups are active. Any node within an active group may participate. The node-count objective penalizes shortfalls relative to the observed active-node targets; counts above those targets incur no penalty. It guides which contacts are selected without enforcing exact count matching.

For each day, contact selection follows four steps.

1. **Form candidate contacts.** In *Operational SBM*, one *proposal round* is one draw of a candidate daily graph from the fitted SBM layer. Repeated rounds enlarge the candidate pool and record how often each node pair is proposed. *Operational Random* instead samples candidate pairs uniformly from nodes in active groups.

2. **Apply participation and turnover rules.** Candidate contacts that start or end in a group not active in the recorded participation sequence are removed. The remaining pairs

are classified as persistent, reactivated, or first-time (new) from the history of the synthetic panel generated up to that day. These are the turnover categories. Churn can be derived and is thus not targeted.

3. **Allocate group-pair quotas.** Within each directed group pair, contacts are selected to approach the observed counts in the three turnover categories. Group-pair and category combinations that contain fewer contacts are handled first. Within a pool, preference is given to pairs with stronger SBM support and to pairs that reduce active-node count shortfalls. SBM support is used only in the SBM-guided regime.

4. **Complete the daily layer.** When a category target cannot be met because its eligible pool is too small, the shortfall is recorded. Remaining candidate capacity is then used to approach the observed total number of contacts for that day.

The three regimes compare complementary information. *Independent SBM* retains learned daily structure without contact-history selection. *Operational Random* applies contact-turnover targets using uniform dyad proposals. *Operational SBM* combines learned trade partner support with contact-history selection. The observed block-participation sequence is preserved across all three regimes. Their comparison assesses learned partner structure and contact-history selection within the preserved block-participation sequence.

After the topology generator selects the directed contacts, a separate statistical model assigns their shipment sizes. We fit this model after the block partition has been inferred. For an edge from node $u$ to node $v$ on day $t$, let $z_u = r$, $z_v = s$, and let $k(u, v) \in \mathcal{K}$ denote the ordered hybrid channel. The generated weight follows

$$w_{uvt} \mid z_u = r, z_v = s, k(u, v) = k, t \sim \mathcal{F}_k(\boldsymbol{\theta}_{rstk}) \, .$$

The distribution family is selected separately for each hybrid channel according to the observed shipment-size scale. Because shipment sizes in the analyzed panels are positive integers, we use a shifted negative-binomial distribution: $x_{uvt} = w_{uvt} - 1$ follows a negative-binomial model with a cell-specific mean and overdispersion. The shift retains the correct minimum value of one animal, while the negative-binomial model allows shipment variability to exceed that of a Poisson model.

The shipment-size model uses the same sender and receiver blocks as the topology and temporal components. For each channel, we estimate shipment-size distributions at six levels: exact day–block-pair–channel, block-pair–channel, day–channel, channel, day, and global. Estimates for sparse, highly specific combinations are pulled toward broader estimates, a form of hierarchical pooling that reduces instability while retaining block- and day-specific information where the data support it. At generation time, the sampler uses the most specific available level in the ordered lookup

$$\text{exact} \rightarrow \text{block-pair} \rightarrow \text{day–channel} \rightarrow \text{channel} \rightarrow \text{day} \rightarrow \text{global.}$$

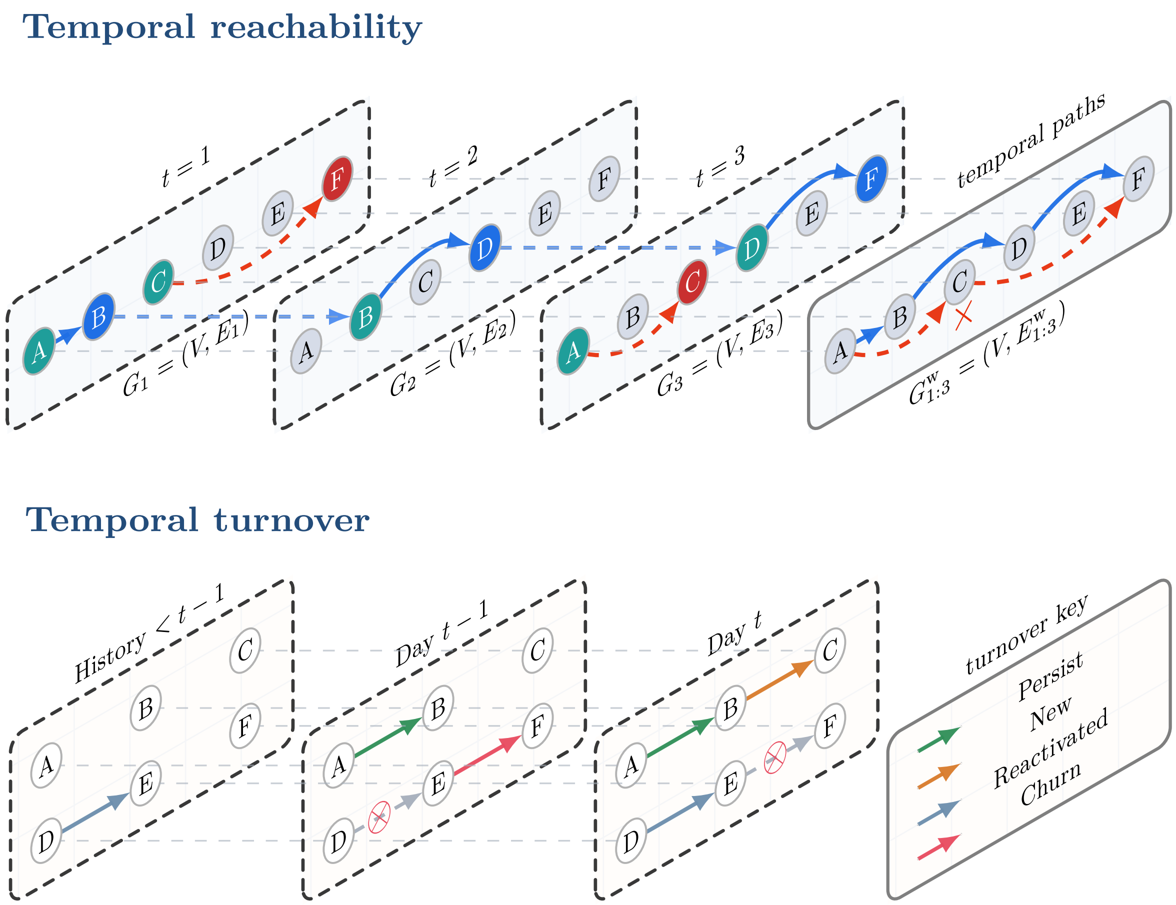


**Supplementary Fig. 3. Temporal reachability and contact turnover categories.** Time-respecting paths may wait across daily layers but cannot use contacts before they occur. Contacts are classified as persistent, new, reactivated, or churned from their occurrence history.

With this approach, contact occurrence and shipment size are modeled separately, but both use the same inferred roles. Weighted block-model research supports the expectation that edge magnitudes vary across latent groups. The separate model used here also keeps the topology ablations interpretable and allows the shipment-size distribution to be changed without refitting the partition.

#### Supplementary Note S5: Structural and transmission validation

Each synthetic panel is compared with the observed panel on the same calendar, node universe, edge direction, and shipment-weight scale. We first confirm quantities that are directly constrained by generation and then test structural outcomes that were not matched explicitly.

Day-level diagnostics include edge and active-node counts, density, reciprocity, observed–synthetic edge and node overlap, movement distance, and shipment-weight summaries. Temporal turnover diagnostics separate first appearances, persistence, reactivation, and churn for both contacts and active nodes. Hybrid-channel diagnostics compare daily contact and weight shares across $F \to F$, $F \to R$, $R \to F$, and $R \to R$. More local checks compare directed group-pair time series and node-level in- and out-activity.

Temporal reachability links these structural checks to possible spreading pathways. A source reaches a target only through contacts that respect the order of the daily layers (Supplementary Fig. 3). In the implementation, each daily layer contributes one propagation step, so multiple contacts cannot be traversed within the same daily snapshot. We summarize the number and fraction of reachable ordered node pairs, newly reachable pairs per day, temporal efficiency, mean arrival time among reached pairs, and source-level forward reach. Further definitions and implementation details are provided in the supplementary online repository.

Structural validation should be accompanied by testing whether observed and generated panels support similar transmission behavior. We therefore apply the same stochastic hybrid disease simulator to the observed panel and to every synthetic panel, using matched initial seed sets and random-number seeds. See Supplementary Table 1

The simulator combines compartmental farm dynamics and regional reservoirs. Regional supernodes do not carry farm compartment states. Instead, each stores a nonnegative and continuous-valued pressure that decays between days. Farm-to-farm movements contribute direct infection hazard, farm-to-region movements seed regional pressure from infectious farms, region-to-farm movements expose susceptible farms to that pressure, and region-to-region movements propagate pressure to receiving regional reservoirs.

**Shipment-weight transformation.** Shipment sizes were transformed for sensitivity analyses that test whether simulated transmission trajectories are sensitive to the edge weights.

For a movement from $u$ to $v$ on day $t$ with shipment size $w_{uvt} > 0$, the transformed contact

**Supplementary Table 1.** Simulation parameters shared by 2018, 2020 and 2022.

| Parameter | Value |
|---|---|
| Transmission: farm → farm | 0.15 |
| Transmission: farm → region | 0.05 |
| Transmission: region → farm | 0.20 |
| Transmission: region → region | 0.04 |
| Daily recovery probability | 0.02 |
| Regional pressure retained each day | 0.90 |
| Background regional pressure per day | 0.0004 |
| Maximum regional pressure | 20 |
| External infection probability per susceptible farm per day | 0.0003 |
| Initially infectious farms | 3 |
| Epidemic replicates per network | 100 |
| Follow-up without contacts | 30 days |

weight was defined as

$$\widetilde{w}_{uvt}^{\text{linear}} = \frac{w_{uvt}}{s_w}$$

under linear weighting, where $s_w > 0$ was estimated once from the observed panel and then held fixed for all observed and generated simulations. Under binary weighting,

$$\widetilde{w}_{uvt}^{\text{binary}} = \mathbb{I}(w_{uvt} > 0) = 1,$$

so shipment size no longer altered the contribution of an observed contact.

**Hybrid transmission process.** The transmission simulation was conducted on a pig-movement system comprising discrete, daily network snapshots, and consisting of a hybrid of farm nodes in a focal region and supernodes representing aggregated dynamics of other regions. This requires separate treatment of different node types.

Let $F$ and $R$ denote the sets of focal farms and regional supernodes, respectively. Let $X_{f,t}$ be the compartmental state of farm $f \in F$ at the start of day $t$, and let $P_{r,t} \geq 0$ be the pressure stored in regional supernode $r \in R$. Initially selected seed farms began in the infectious state, all other farms began susceptible, and regional pressure was initialized at zero. Regional supernodes did not carry susceptible, infectious, or recovered states.

Let $\widetilde{w}_{uvt}$ denote the transformed and scaled weight of directed contact $(u, v) \in E_t$. Before entering the transmission equations, regional pressure was bounded as

$$\overline{P}_{r,t} = \min(P_{r,t}, P_{\max}),$$

where $P_{\max}$ is the maximum permitted reservoir pressure. For a susceptible farm $f$, the daily hazard was the sum of direct farm-to-farm transmission and exposure through incoming region-to-farm movements:

$$h_{f,t} = h_{f,t}^{F\to F} + h_{f,t}^{R\to F} + h_0,$$

where

$$h_{f,t}^{F\to F} = \beta_{FF}\,\chi\,\iota \sum_{\substack{u\in F\\(u,f)\in E_t}} \widetilde{w}_{uft}\,\mathbf{1}\{X_{u,t}=I\}\,,$$

and

$$h_{f,t}^{R\to F} = \beta_{RF}\,\chi \sum_{\substack{r\in R\\(r,f)\in E_t}} \widetilde{w}_{rft}\,\log\bigl(1+\overline{P}_{r,t}\bigr)\,.$$

Here, $\beta_{FF}$ and $\beta_{RF}$ are channel-specific transmission coefficients, while $\iota$ and $\chi$ are the farm infectiousness and susceptibility multipliers. Throughout our study, $\iota$ and $\chi$ were always set to 1.0 and kept constant. The term $h_0$ represents an exogenous farm-level infection hazard and is zero when exogenous infection is disabled. Conditional on being susceptible at the start of the day, farm $f$ became infected with probability

$$\Pr(X_{f,t+1}=I \mid X_{f,t}=S) = 1-\exp[-\min(h_{f,t},h_{\max})]\,,$$

where $h_{\max}$ bounds the numerical hazard.

Regional pressure was updated through persistence of existing pressure, export from infectious farms, and circulation among regional supernodes:

$$P_{r,t+1} = \left[\rho P_{r,t} + \beta_{FR}\,\iota \sum_{\substack{u\in F\\(u,r)\in E_t}} \widetilde{w}_{urt}\,\mathbf{1}\{X_{u,t}=I\} + \beta_{RR} \sum_{\substack{q\in R\\(q,r)\in E_t}} \widetilde{w}_{qrt}\,\log\bigl(1+\overline{P}_{q,t}\bigr)\right]_0^{P_{\max}}\,,$$

where $\rho$ is the daily reservoir-retention factor, $\beta_{FR}$ controls pressure exported from infectious farms, $\beta_{RR}$ controls pressure transferred between regional supernodes, and

$$[x]_0^{P_{\max}} = \min\{P_{\max},\max(0,x)\}\,.$$

Under SIS dynamics, a farm that was infectious at the start of day $t$ recovered to the susceptible state with probability $\gamma$ and otherwise remained infectious. Under SIR dynamics, recovery occurred with the same probability but led to an absorbing recovered state. All infection hazards and regional-pressure contributions for day $t$ were calculated from $X_{\cdot,t}$ and $P_{\cdot,t}$, after which farm states and regional pressures were updated synchronously. Consequently, a farm infected on day $t$ could not transmit until day $t+1$, and pressure added to a regional supernode on day $t$ could not expose farms until a later daily layer. Daily incidence was the number of new farm infections generated during this update, whereas daily prevalence was the number of farms infectious after the update.

After the final movement layer, the simulation continued for $T_{\text{tail}}$ contact-free days to allow farm infections and regional pressure to resolve. Daily trajectory summaries were calculated over the movement-panel calendar, whereas full-horizon outcomes, including whether a farm was ever infected, used the complete simulation horizon.

**Transmission scenarios and sensitivity analyses.** The main analysis combined SIS farm dynamics with linear shipment weighting. Infectious farms recovered to the susceptible state with the specified daily recovery probability and could subsequently be reinfected. We used two sensitivity scenarios:

1. **SIR with linear shipment weights.** The linear contact-weight transformation was retained, but recovered farms entered a removed state and could not be reinfected.
2. **SIS with binary contact weights.** The SIS transition rules were retained, but every positive movement received unit weight regardless of shipment size.

The same observed panel, generated panels, transmission coefficients, recovery probability, regional-pressure parameters, initial farm seed sets, random-number seeds, simulation horizon, and replicate design were used across scenarios. The sensitivity analyses therefore changed only the farm-state model or the shipment-weight representation relative to the main SIS–linear setting. See Supplementary Table 2

**Supplementary Table 2.** Main and sensitivity analyses. All other parameters were shared.

| Analysis | Farm dynamics | Contact weight |
|---|---|---|
| Main | SIS | Shipment size |
| Sensitivity 1 | SIR | Shipment size |
| Sensitivity 2 | SIS | 1 per contact |

**Transmission outcomes and aggregation.** In the reported analysis, each generation regime was represented by 10 independently generated panels, each evaluated with 10 epidemic replicates. The observed panel was evaluated once with the same 10 matched initial-seed sets and random-number seeds, which were reused for every generated panel.

Let $\mathcal{F}$ denote the set of focal farms and let $N_F = |\mathcal{F}|$. For panel $p$ and epidemic replicate $r$, let $Y_{fpr} = 1$ if focal farm $f$ was infected at least once during the full simulation horizon, including infection at initialization, and let $Y_{fpr} = 0$ otherwise. The replicate-specific farm attack rate was defined as

$$A_{pr} = \frac{1}{N_F} \sum_{f \in \mathcal{F}} Y_{fpr}.$$

Each farm therefore contributed at most once to the attack rate, regardless of recovery and reinfection under SIS dynamics, and farms selected as initial seeds were included.

Let $I_{fprt} = 1$ if focal farm $f$ was infectious on panel day $t$, and let $I_{fprt} = 0$ otherwise. The replicate-specific farm peak prevalence was defined as

$$M_{pr} = \max_{t \in \mathcal{T}} \sum_{f \in \mathcal{F}} I_{fprt}.$$

This quantity is the largest daily number of simultaneously infectious focal farms. It was calculated separately within each replicate before aggregation and was not obtained by taking the maximum of the replicate-mean prevalence curve.

For a generation regime containing $P$ independently generated panels and $R$ epidemic replicates per panel, the reported mean attack rate and mean peak prevalence were

$$\overline{A}_{\text{gen}} = \frac{1}{PR}\sum_{p=1}^{P}\sum_{r=1}^{R} A_{pr}, \qquad \overline{M}_{\text{gen}} = \frac{1}{PR}\sum_{p=1}^{P}\sum_{r=1}^{R} M_{pr}.$$

The corresponding observed-panel means were calculated across the $R$ matched epidemic replicates:

$$\overline{A}_{\text{obs}} = \frac{1}{R}\sum_{r=1}^{R} A_{0r}, \qquad \overline{M}_{\text{obs}} = \frac{1}{R}\sum_{r=1}^{R} M_{0r},$$

where $p = 0$ denotes the observed panel. Because every generated panel used the same number of matched replicates, averaging the panel-specific means is equivalent to pooling all $PR$ replicate-specific outcomes. The scalar-outcome distributions for a generated regime likewise pool the $PR$ replicate-specific values across its generated panels.

For each focal farm $f$, network-generated attack probability was calculated conditional on that farm not being selected as an initial seed. Let $S_{fr} = 1$ when farm $f$ belonged to the initial-seed set for matched replicate $r$, and let $S_{fr} = 0$ otherwise. Within generated panel $p$, the probability was estimated as

$$\pi_{fp} = \frac{\sum_{r=1}^{R}(1 - S_{fr})Y_{fpr}}{\sum_{r=1}^{R}(1 - S_{fr})}.$$

Replicates in which farm $f$ was initially seeded were therefore excluded from both the numerator and denominator. For a generation regime, the farm-level estimate was averaged across its generated panels,

$$\overline{\pi}_{f,\text{gen}} = \frac{1}{P}\sum_{p=1}^{P} \pi_{fp},$$

whereas the observed-panel estimate was $\pi_{f0}$. Because the same matched seed sets were reused for every generated panel, this panel-averaged estimate is equivalent to pooling all eligible panel–replicate combinations. Farm-level map differences were calculated as $\overline{\pi}_{f,\text{gen}} - \pi_{f0}$.

Other evaluated outcomes included daily infectious-farm prevalence, farm incidence, cumulative incidence, peak timing, outbreak duration, prevalence and reservoir areas under the curve, regional reservoir summaries, channel-specific hazard contributions, and regional spatial agreement.

### Supplementary Results

In this section we report the results to further support the findings in the main study. This includes the observed (2018 panel) network and block structure, the results of the sensitivity analyses, and the additional validation results on the 2020 and 2022 panels.

#### Supplementary Note S6: Observed hybrid-network structure

The 2018 CR35 panel combines very sparse daily contact with substantial movement through regional supernodes. This suggested that the network generators must allocate a small number of daily contacts across hybrid channels with very different numbers of possible pairs, while preserving weekly activity and external movement pressure. Supplementary Fig. 4 and Supplementary Table 3 summarize these constraints.

The panel contains 843 explicit focal farms and 38 regional supernodes over 365 daily layers. These 881 nodes define 775,280 possible directed non-self pairs per day. However, even the busiest day contains fewer than 500 contacts. Daily all-pair occupancy remains below 0.1%.

**Supplementary Table 3.** Post-aggregation annual contacts and movement weight by hybrid channel. Directed contacts are daily ordered pair contacts after repeated movements on the same day are summed into one weighted contact. Distinct directed pairs count ordered node pairs observed at least once during 2018.

| | Annual directed contacts | | Distinct directed pairs | | Movement weight | |
|---|---|---|---|---|---|---|
| Channel | Count | Share | Count | Share | Total | Share |
| F→F | 15,110 | 19.0% | 1,386 | 32.0% | 2,325,826 | 9.4% |
| F→R | 22,725 | 28.5% | 1,719 | 39.7% | 3,604,368 | 14.6% |
| R→F | 10,237 | 12.9% | 609 | 14.1% | 4,077,495 | 16.5% |
| R→R | 31,592 | 39.7% | 615 | 14.2% | 14,651,061 | 59.4% |
| **Total** | **79,664** | **100.0%** | **4,329** | **100.0%** | **24,658,750** | **100.0%** |

Across the year, the panel contains 79,664 daily directed contacts on 4,329 distinct directed pairs, with total movement weight 24,658,750. Farm–farm contacts account for 32.0% of distinct pairs but only 19.0% of daily contacts and 9.4% of movement weight. The three channels involving regional supernodes contribute 81.0% of contacts and 90.6% of weight. Region–region movements are especially concentrated: 14.2% of distinct pairs carry 39.7% of contacts and 59.4% of total weight. The regional nodes therefore represent a dominant component of the hybrid network, although region–region movements do not directly contact focal farms.

Daily activity also follows a strong calendar pattern (see Supplementary Fig. 4, outer rings, follow the clockwise direction for the daily trade activities). Contact and active-node counts (Supplementary Fig. 4, outermost ring, stacked bars) fall on weekends and Dutch public holidays (as indicated by the light-blue and red backgrounds) and recover on weekdays (white background), while fewer than roughly 30% of nodes are active on a typical plotted day. The observed panel is thus sparse, directed, weighted, calendar-structured, and regionally coupled. These are the baseline properties against which the generated panels are evaluated.

The central network in Supplementary Fig. 4 visualizes the aggregated trade activities between farms and regions in 2018, organized by the inferred network block structure. The dendrogram on top of the central network visualizes how the farms and regions were grouped into 11 leaf blocks and then further merged into higher-level blocks. Note that the central network is not related to the calendar dates shown in the outer visualizations.

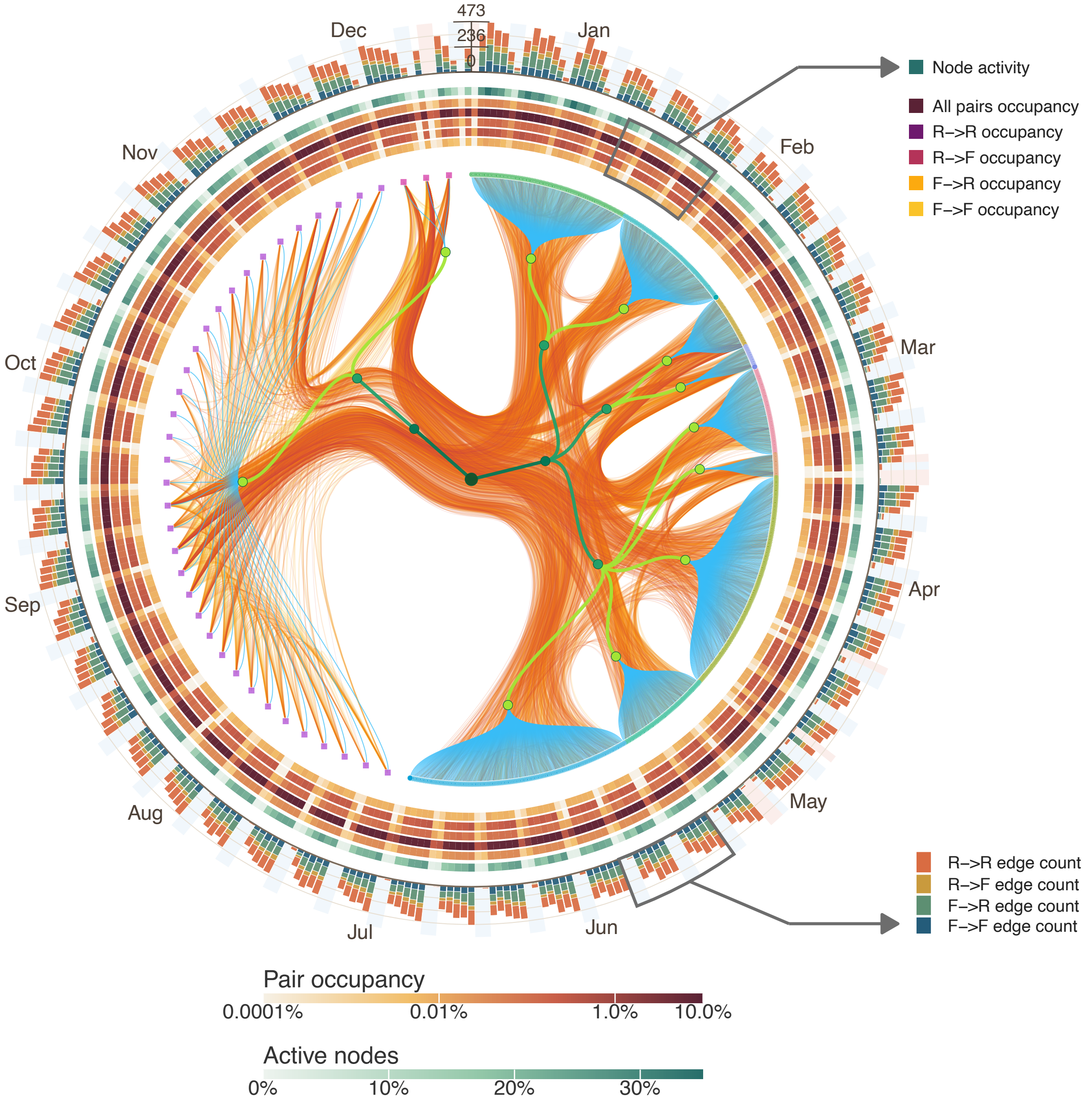


**Supplementary Fig. 4. Daily activity and inferred nested structure of the 2018 hybrid network.** The central network is arranged by the inferred hierarchical block structure and distinguishes observed farm nodes (circles) and regional supernodes (squares). The edges in the bottom layer represent trade activities aggregated across 2018 with color indicating log-transformed shipment sizes. The dendrogram on top visualizes how the farms and regions share the same grouping into leaf blocks and higher-level blocks. Surrounding rings show daily pair occupancy by hybrid channel, and outer bars show daily directed contact counts, with weekends and Dutch public holidays marked by light-blue and red backgrounds respectively.

#### Supplementary Note S7: Detailed block roles and metadata interpretation

Metadata substantially changes the resolution of the inferred network roles in the observed pig-movement network. Without metadata, the fitted data nodes collapse to one farm block and one regional block. With metadata annotations, the sampled joint model resolves 11 leaf blocks, including nine farm blocks and two regional blocks. The additional farm blocks reveal heterogeneity that is only weakly supported by the sparse daily movement layers alone.

We distinguish inferred network roles from registry categories in the metadata. In our definition, a source block mainly sends animals, a sink block mainly receives them, a mixed trading block both sends and receives substantial movement, and an external-pressure block contains only regional supernodes. Meanwhile, breeding, nursery, finisher, and other labels are observed production-stage metadata. A production stage may occupy different network roles depending on local trading and exchange with the external system.

For each leaf block, we calculate the share of incident movement weight that is outgoing,

$$p_{\text{out}}(b) = \frac{\sum_j W_{bj}}{\sum_j W_{bj} + \sum_i W_{ib}}.$$

Farm blocks with $p_{\text{out}} \geq 0.70$ are classified as sources, those with $p_{\text{out}} \leq 0.30$ as sinks, and the remainder as mixed trading blocks. The rule identifies four farm source blocks, four mixed trading blocks, one sink block, and two regional external-pressure blocks from the 11 leaf blocks (Supplementary Table 4).

| Block | Class | Out | In | Out share | Role |
|---|---|---|---|---|---|
| B261 | farm | 1,429,163 | 320,962 | 0.817 | source |
| B391 | farm | 1,013,679 | 254,162 | 0.800 | source |
| B452 | farm | 662,241 | 276,829 | 0.705 | source |
| B231 | farm | 404,368 | 144,474 | 0.737 | source |
| B190 | farm | 590,438 | 527,073 | 0.528 | mixed trading |
| B187 | farm | 419,412 | 594,768 | 0.414 | mixed trading |
| B283 | farm | 607,496 | 378,430 | 0.616 | mixed trading |
| B62 | farm | 568,031 | 316,610 | 0.642 | mixed trading |
| B646 | farm | 235,366 | 3,590,013 | 0.062 | sink |
| B856 | regional | 11,773,817 | 12,675,348 | 0.482 | external pressure |
| B878 | regional | 6,954,739 | 5,580,081 | 0.555 | external pressure |

**Supplementary Table 4.** *Leaf-block role assignment from directed in–out balance.* Block IDs are arbitrarily generated during block inference. Out and In are total directed movement weights leaving and entering each block. Out share is $p_{\text{out}}$. Farm roles use the 70% source and 30% sink cutoffs; regional supernode blocks are treated as external pressure roles.

The directed block-mixing profiles provide a different view of these roles. In Supplementary Fig. 5, left panel, animal movement weights (shipment sizes) mainly went through regional (external) pathways, including region–region circulation and movements from regional blocks into the farm sink. This is reflected by the overall higher block-pair trade intensities as represented by the heights (and colors) of the cubes.

In Supplementary Fig. 5, right panel, after each source block is normalized by its outgoing total, the receiving profiles (columns that represent receiver blocks) separate the same four source blocks, four mixed blocks, one sink, and two regional blocks more clearly. Agreement between the profile clustering and the in–out balance shows that our initial role assignments can reflect the organization of the movement weights in the observed trade network.

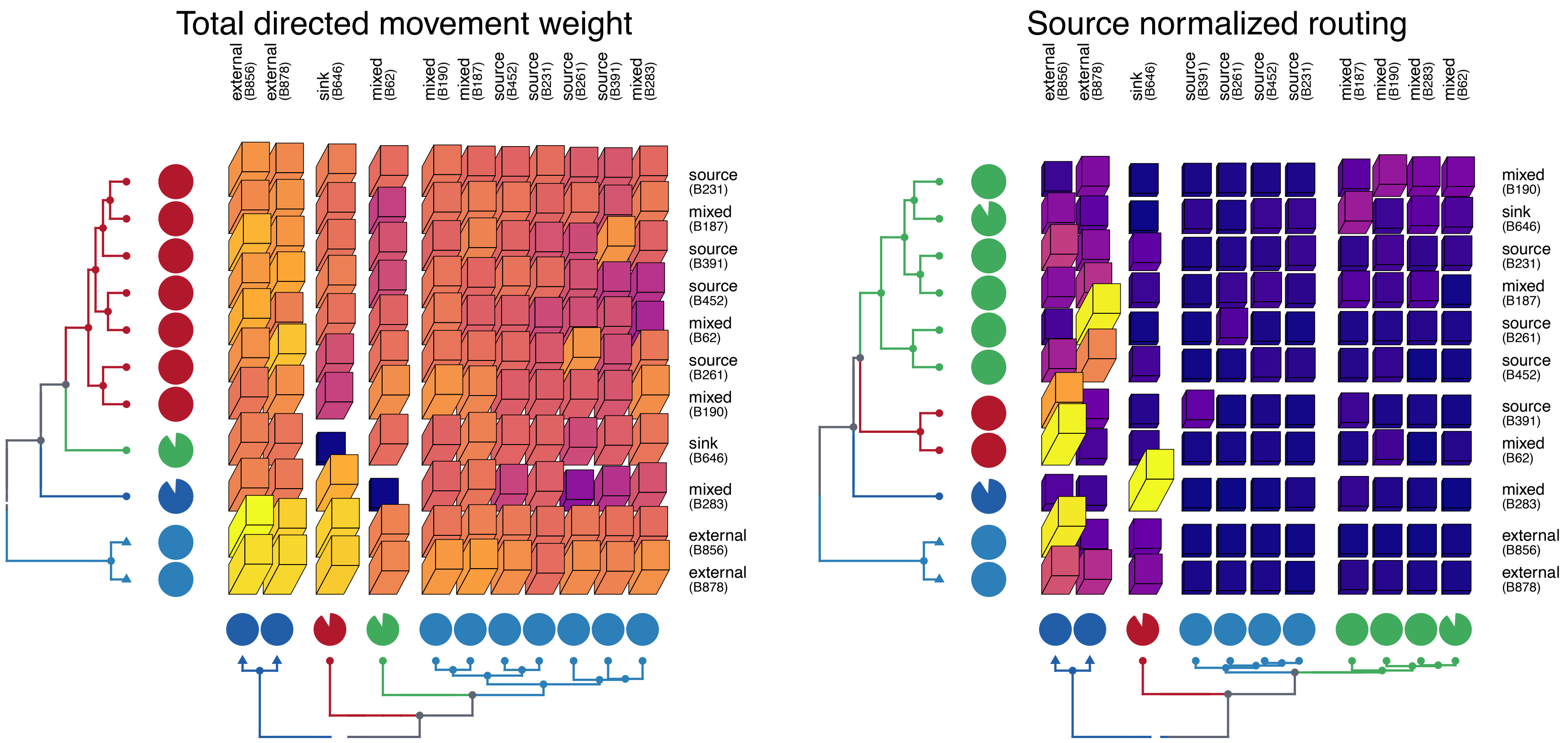


**Supplementary Fig. 5. Directed block mixing matrix among joint metadata leaf blocks.** Each cube in the matrix represents a directed block-pair. Their heights and colors indicate the trade intensities between the directed block-pairs. Rows represent sender blocks and columns represent receiver blocks. The left heatmap shows log-transformed observed movement weight; the right heatmap normalizes each row by the source block's total outgoing weight. The pie beside each row shows the proportion of target blocks receiving nonzero movement from that source block; the pie below each column shows the proportion of source blocks contributing nonzero movement to that target block. The pies measure the breadth of outgoing and incoming connections. Pie colors match the corresponding dendrogram clusters. Row dendrograms group blocks by their outgoing profiles, while column dendrograms group blocks by their incoming profiles, using a node-class constraint that keeps regional blocks distinct from farm blocks. Triangular tips denote regional supernode blocks and circular tips denote farm blocks.

#### Supplementary Note S8: Sensitivity of transmission results to farm-state dynamics and shipment weighting

We tested whether the transmission comparison depended on two choices in the hybrid simulator: whether recovered farms returned to susceptibility and whether shipment size affected the contact hazard. The observed panel and the same generated panels were rerun using the matched seed sets, random-number seeds, epidemiological parameters, and replicate design of the main analysis. The identity of the best-performing regime remained unchanged in both sensitivity settings: *Operational SBM* produced the closest agreement with the observed-panel simulations, while *Independent SBM* and *Operational Random* showed larger trajectory and burden discrepancies.

**SIR dynamics with linear shipment weights.** Replacing SIS with SIR dynamics reduced the absolute differences between the observed and generated prevalence trajectories. Because recovered farms could not return to the susceptible state, prevalence no longer approached the sustained levels seen under SIS dynamics. For *Operational SBM*, the remaining discrepancy was concentrated near the beginning of the simulation, when its generated prevalence trajectory was slightly below the observed-panel trajectory. Despite this early underestimation, *Operational SBM* remained closer to the observed panel than either ablation in trajectory shape, epidemic magnitude, and farm-level burden (Supplementary Fig. 6).

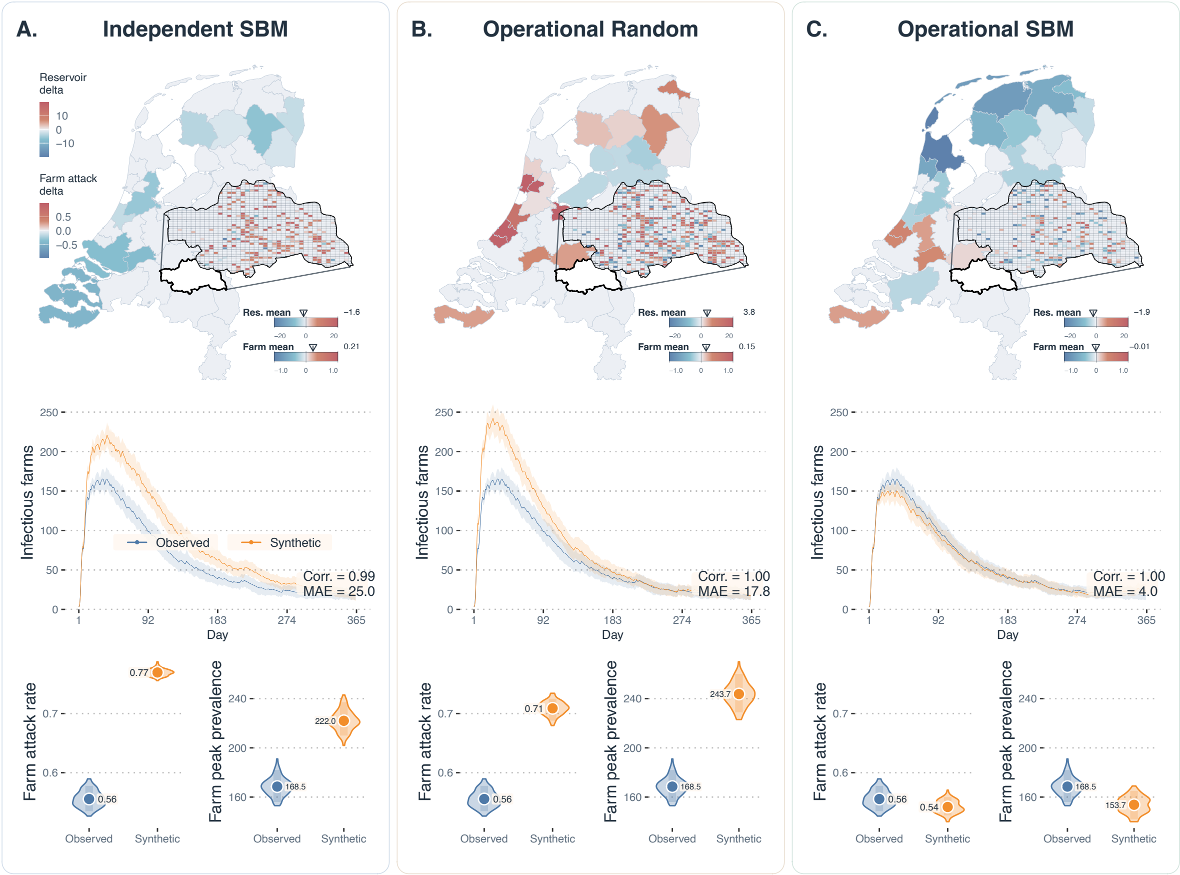


**Supplementary Fig. 6. Sensitivity of transmission results to SIR farm dynamics with linear shipment weights.** Farm nodes follow an SIR process in which recovered farms enter a removed state and cannot be reinfected; regional supernodes retain the same movement-mediated pressure process as in the main analysis. Shipment sizes contribute linearly to transmission hazards after scaling by the fixed observed-panel weight scale. Each column compares one generation regime with the observed panel using the same generated panels, epidemiological parameters, initial seed sets, random-number seeds, and replicate design as the main analysis. Top-row panels show regional reservoir-pressure and farm-level attack-probability differences. Middle-row panels compare mean daily infectious-farm prevalence and replicate intervals. Bottom-row panels compare farm attack rate and peak prevalence. Absolute trajectory differences are smaller than under the main SIS setting, with a slight early underestimation of prevalence, under *Operational SBM*, which retains the closest overall agreement.

**SIS dynamics with binary contact weights.** Replacing linear shipment weighting with binary contact weights also reduced the absolute differences between observed and generated prevalence trajectories for the two ablation regimes, although the smaller *Operational SBM* gap increased modestly. Treating every positive shipment as a unit contact removed variation in transmission pressure arising from shipment size and therefore reduced the influence of differences in generated movement weights. Some overestimation of prevalence nevertheless remained, showing that shipment-size variation did not fully account for the transmission discrepancies. *Operational SBM* again gave the closest agreement with the observed panel, while the two ablation regimes retained larger deviations (Supplementary Fig. 7).

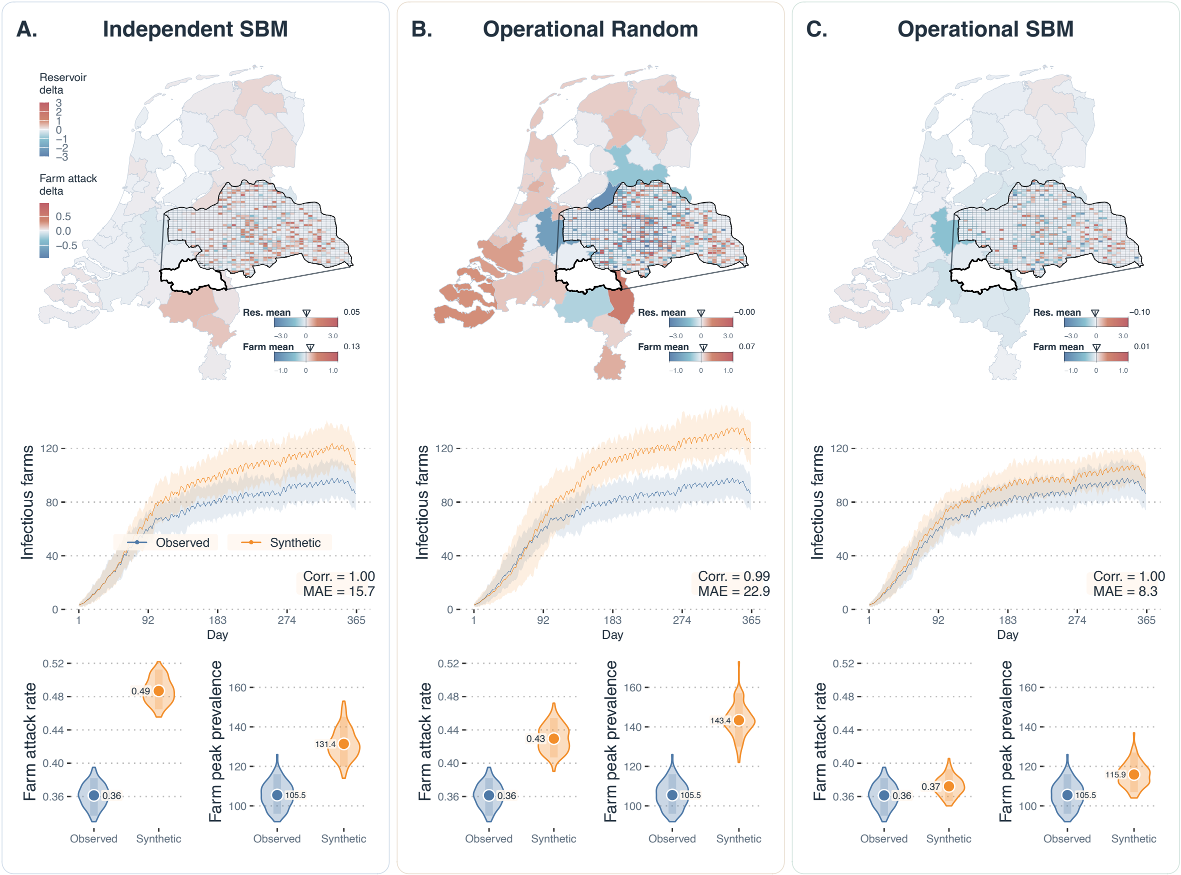


**Supplementary Fig. 7. Sensitivity of transmission results to binary contact weights under SIS dynamics.** Farm nodes follow the SIS process used in the main analysis, but every positive movement contributes unit contact weight regardless of shipment size. Regional supernodes retain the same movement-mediated pressure process. Each column compares one generation regime with the observed panel using the same generated panels, epidemiological parameters, initial seed sets, random-number seeds, and replicate design as the main analysis. Top-row panels show regional reservoir-pressure and farm-level attack-probability differences. Middle-row panels compare mean daily infectious-farm prevalence and replicate intervals. Bottom-row panels compare farm attack rate and peak prevalence. Binary weighting markedly narrows the observed–generated trajectory gaps of the two ablation regimes, although some overestimation of prevalence remains. *Operational SBM* retains the closest agreement with the observed-panel simulations.

Across the two sensitivity settings, changes to the epidemic process and shipment weighting altered the absolute scale and timing of the observed–generated differences but retained *Operational SBM* as the closest regime. The smaller gaps under binary weighting indicate that shipment-size scaling contributes to the quantitative transmission discrepancy. The residual overestimation under binary weighting, together with the continued advantage of *Operational SBM* under SIR dynamics, shows that differences in network topology and temporal contact organization remain relevant when shipment-size effects or reinfection are modified. These analyses support the main structural conclusion while showing that the precise magnitude of epidemic disagreement is conditional on the downstream transmission model.

#### Supplementary Note S9: Additional results on the 2020 and 2022 panels

**Additional validation using the 2020 panel.** We applied the same block inference, turnover and reachability, and SIS transmission analyses used for the 2018 evaluation to the complete 2020 panel. Results are shown below.

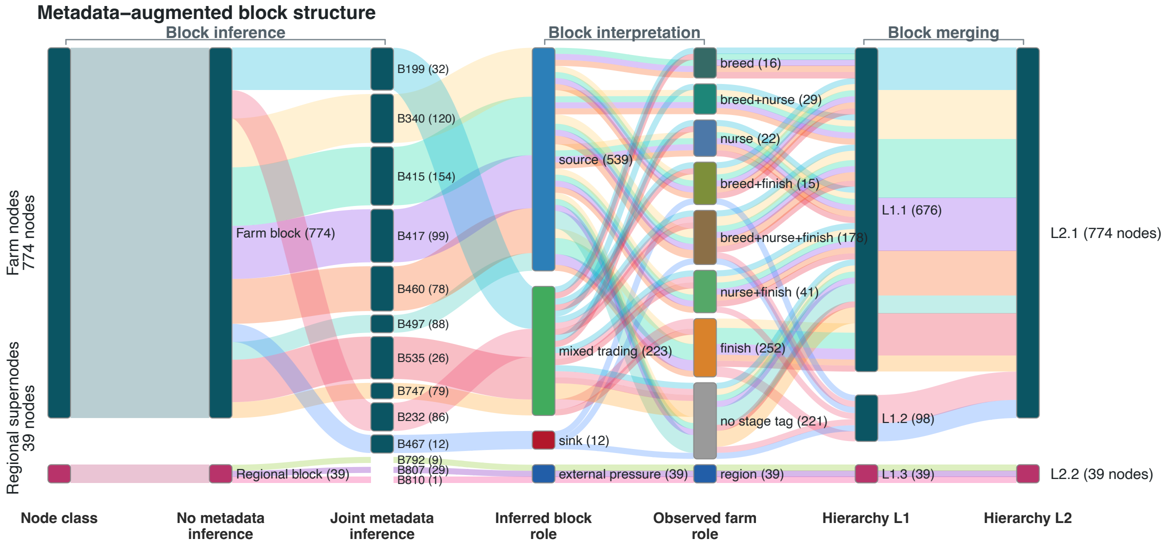


**Supplementary Fig. 8. Additional temporal validation: metadata and block inference in the 2020 panel.** The Sankey traces nodes from their initial classes through the no-metadata and joint-metadata partitions, inferred movement roles, observed production stages, and successive hierarchy levels. Regional supernodes are displayed separately from farm blocks.

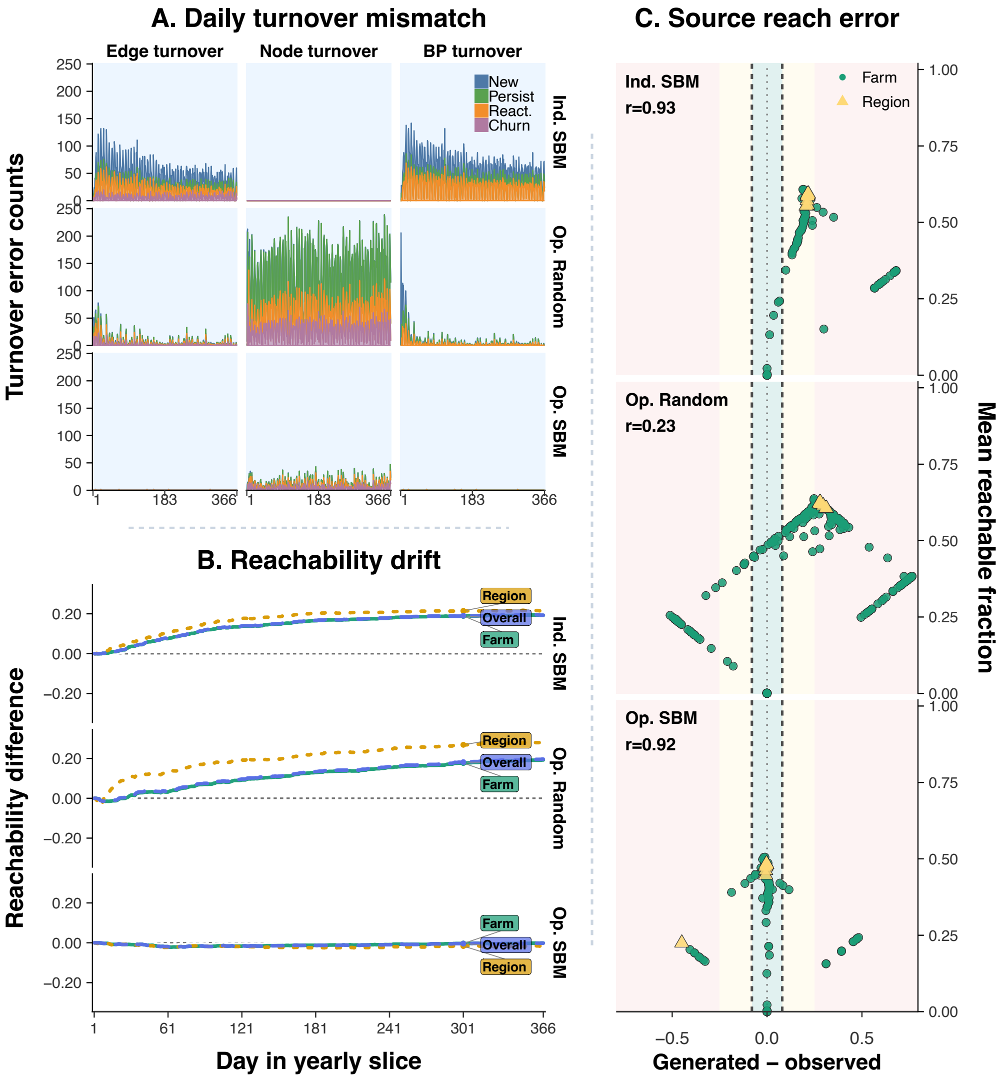


**Supplementary Fig. 9. Additional temporal validation: turnover and temporal reachability in the 2020 panel. A**: Daily absolute generated-versus-observed differences in edge turnover, active-node turnover, and contact turnover within directed block pairs (BP turnover) under each generative regime. Colors distinguish new, persistent, reactivated, and, for edges and nodes, churn events. **B**: Generated-minus-observed temporal reachability for farm sources, regional sources, and all sources. Reachability is the fraction of eligible ordered source–target pairs connected by a chronologically ordered directed path in the contacts accumulated through each day. **C**: Source-level final forward-reach error under each generative regime. The horizontal axis shows generated minus observed reach, and the vertical axis shows their mean. Shading denotes absolute-error ranges, and annotations report the Spearman rank correlation between observed and generated source-level reach.

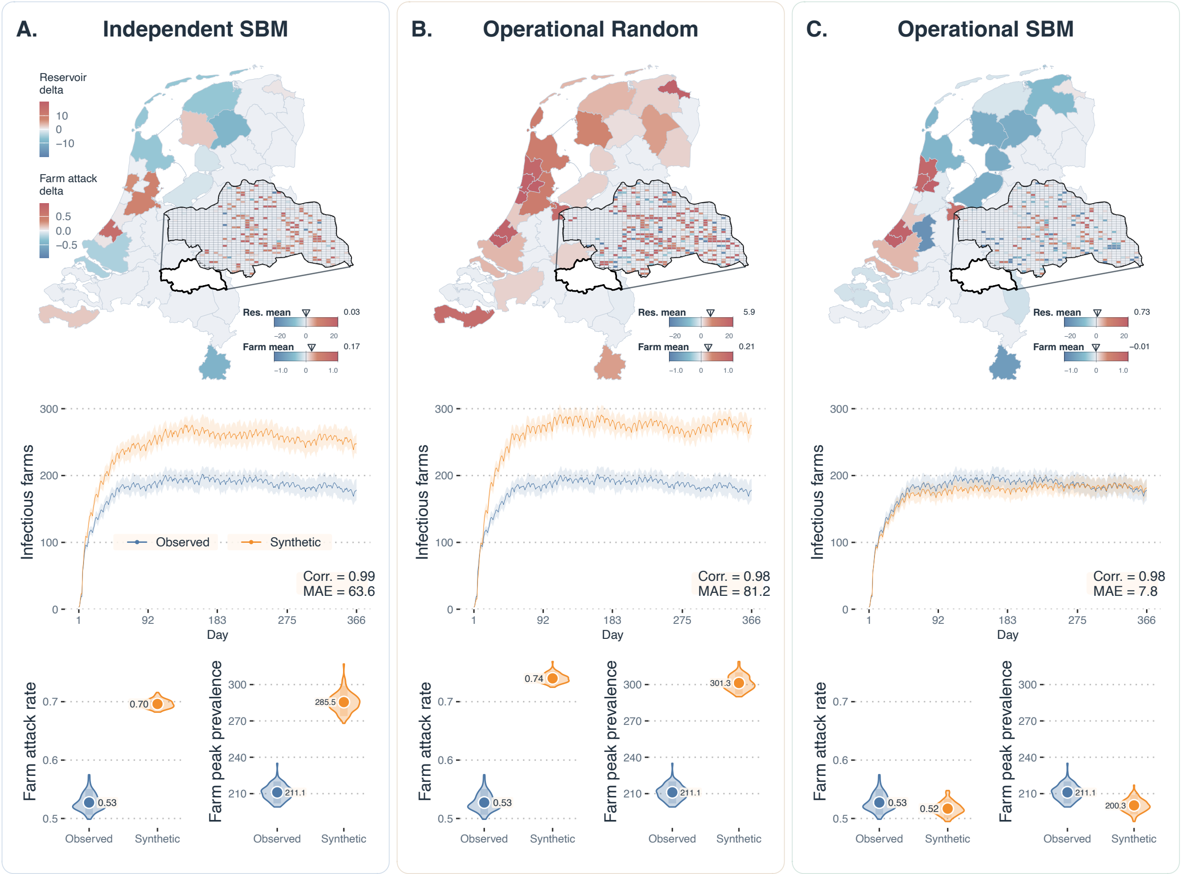


**Supplementary Fig. 10. Additional temporal validation: SIS transmission simulations in the 2020 panel.** Farm nodes follow an SIS process, while regional supernodes carry continuous movement-mediated pressure. Each column compares a generative regime with the observed panel using matched simulation parameters, initial seeding farms, and random-number seeds. The maps show final-day generated-minus-observed regional reservoir pressure and spatially aggregated farm attack-probability differences within CR35. The middle panels show the mean daily number of infectious farms with replicate percentile bands. The bottom panels show replicate distributions of farm attack rate and peak infectious-farm count.

**Additional validation using the 2022 panel.** We applied the same block inference, turnover and reachability, and SIS transmission analyses used for the 2018 evaluation to the complete 2022 panel. Results are shown below.

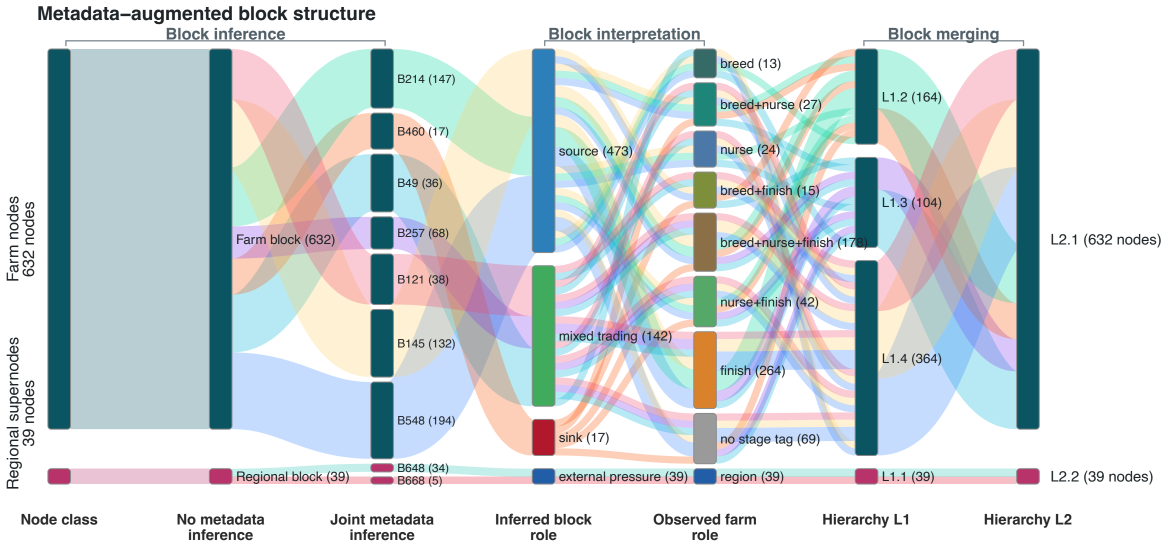


**Supplementary Fig. 11. Additional temporal validation: metadata and block inference in the 2022 panel.** The Sankey traces nodes from their initial classes through the no-metadata and joint-metadata partitions, inferred movement roles, observed production stages, and successive hierarchy levels. Regional supernodes are displayed separately from farm blocks.

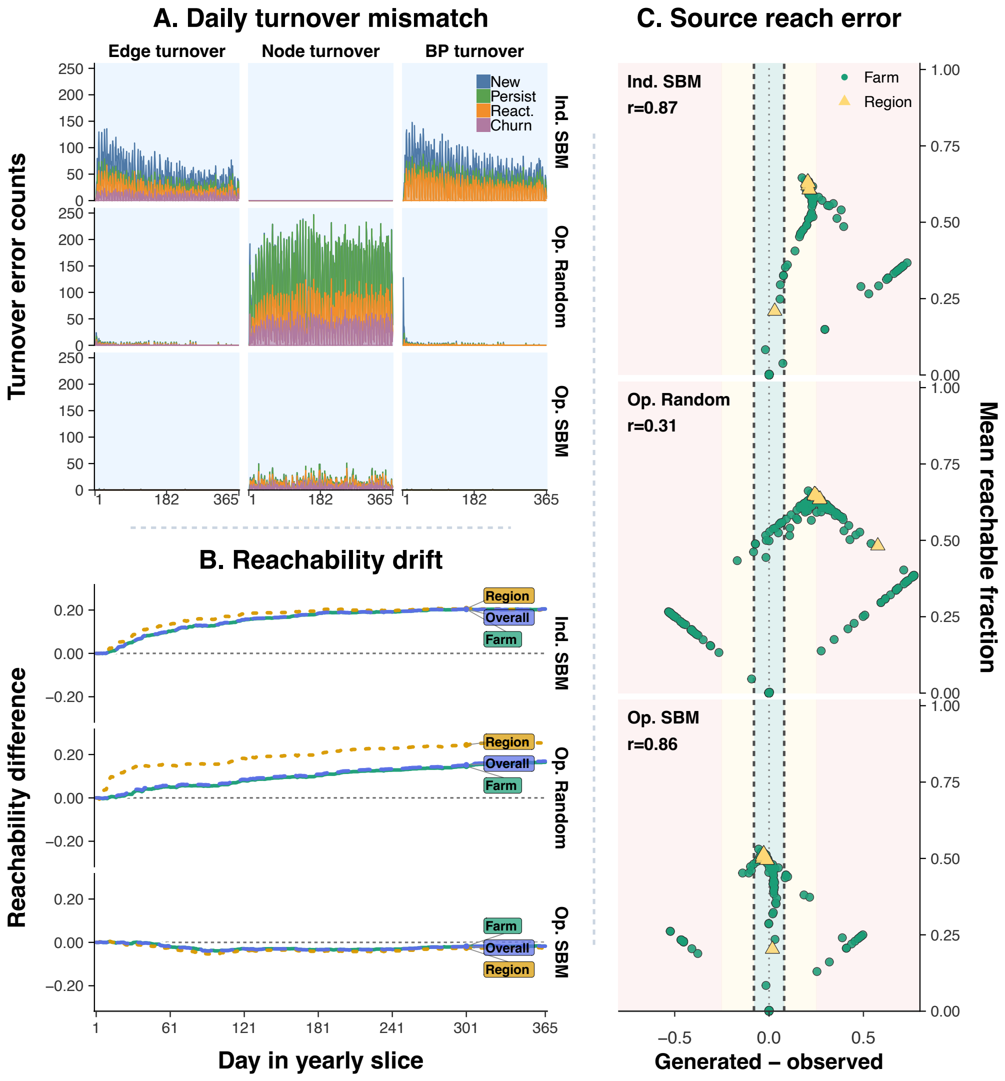


**Supplementary Fig. 12. Additional temporal validation: turnover and temporal reachability in the 2022 panel. A**: Daily absolute generated-versus-observed differences in edge turnover, active-node turnover, and contact turnover within directed block pairs (BP turnover) under each generative regime. Colors distinguish new, persistent, reactivated, and, for edges and nodes, churn events. **B**: Generated-minus-observed temporal reachability for farm sources, regional sources, and all sources. Reachability is the fraction of eligible ordered source–target pairs connected by a chronologically ordered directed path in the contacts accumulated through each day. **C**: Source-level final forward-reach error under each generative regime. The horizontal axis shows generated minus observed reach, and the vertical axis shows their mean. Shading denotes absolute-error ranges, and annotations report the Spearman rank correlation between observed and generated source-level reach.

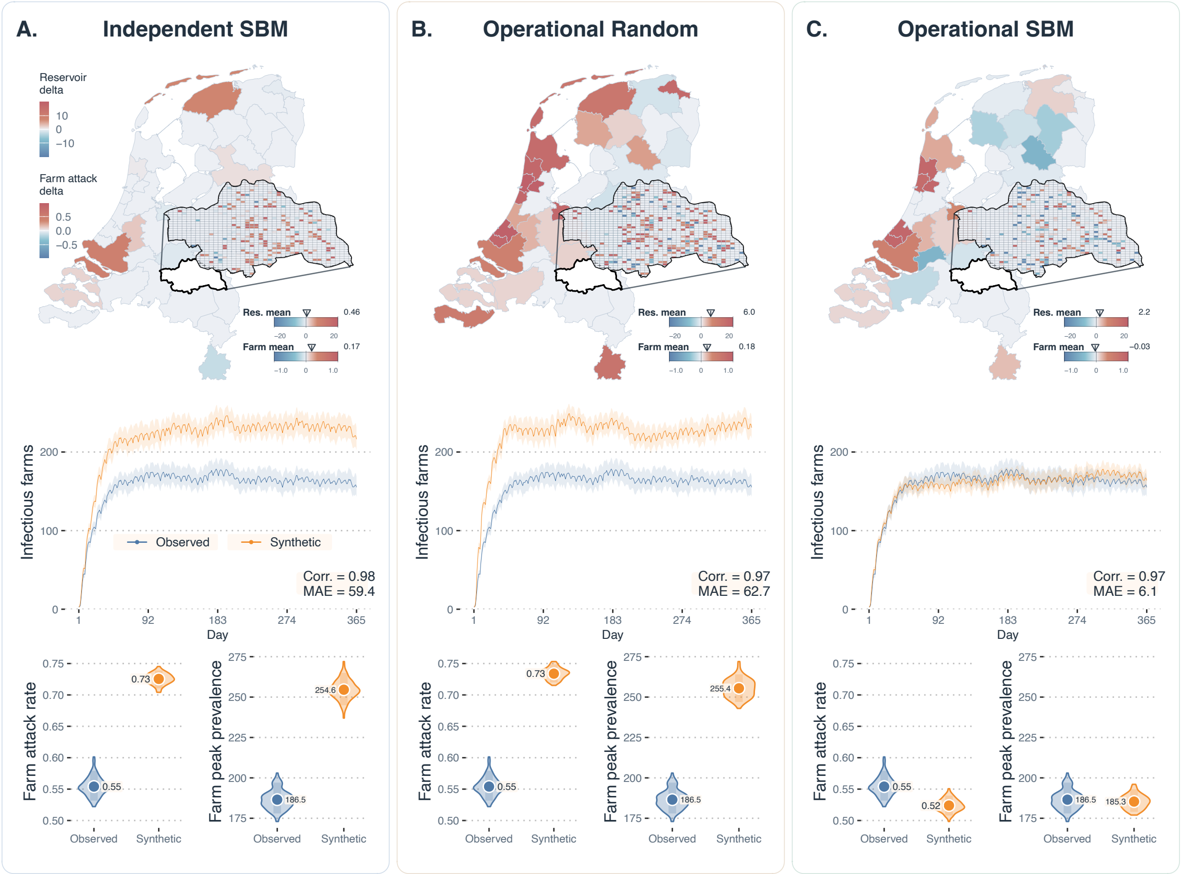


**Supplementary Fig. 13. Additional temporal validation: SIS transmission simulations in the 2022 panel.** Farm nodes follow an SIS process, while regional supernodes carry continuous movement-mediated pressure. Each column compares a generative regime with the observed panel using matched simulation parameters, initial seeding farms, and random-number seeds. The maps show final-day generated-minus-observed regional reservoir pressure and spatially aggregated farm attack-probability differences within CR35. The middle panels show the mean daily number of infectious farms with replicate percentile bands. The bottom panels show replicate distributions of farm attack rate and peak infectious-farm count.

#### Supplementary Analysis

In this analysis, we connect the evaluation of network fidelity with the discussion of disclosure risk. Using GIAB pig-stock data, we first compare stock–activity associations and aggregate-degree rank patterns in the observed and synthetic panels. These comparisons assess how the operational and SBM components contribute to preserving holding-level activity and distinct trade partner structure, providing a supplementary structural check alongside the temporal-reachability analysis. We then test whether pig-stock ranks can identify anonymous farm nodes through their activity ranks under two candidate populations. Comparing linkage success across the observed network and all three generation regimes quantifies how synthesis changes disclosure risk under this specific ranking approach.

##### Supplementary Note S10: Pig-stock ranks and anonymous network activity

**Comparison population and metrics.** In the heatmaps, we compare GIAB pig-stock ranks with activity ranks in the observed 2018 CR35 panel and the three synthetic versions. Pig totals were summed across pig-category records for each holding identifier. 569 holdings with positive stocks were ranked in every panel; 274 of the 843 holdings had no pig record and were excluded. The heatmaps help to assess the retention of pig-stock-related structure after generation.

Total degree is computed by summing incoming and outgoing directed daily contacts; total weighted degree is equivalent to summing shipment sizes; and aggregate degree counts distinct incoming and outgoing partners over the year. Intuitively, repeated contacts increase total degree of an animal holding, but only count once per direction towards aggregate degree. A pair of holdings trading in both directions contributes two. Pig totals and all the network metrics were ranked in descending order with average ranks for ties.

**Retained stock–activity associations.** Observed pig stock correlated strongly with total degree ($\rho = 0.705$), total weighted degree (0.686), and less strongly with aggregate degree (0.357). In the same metric order, correlations were 0.705, 0.671, and 0.500 for *Independent SBM*; 0.226, 0.219, and 0.090 for *Operational Random*; and 0.568, 0.550, and 0.436 for *Operational SBM* (Supplementary Fig. 14). *Independent SBM* matched all 843 holdings' annual total degrees exactly. Its stock–weighted-degree correlation nevertheless differed by 0.015. *Operational SBM* recovered much of the association lost under uniform pairing and was closest for the stock–aggregate-degree association.

**Partner-count bands.** The observed panel and both operational regimes showed pronounced horizontal bands in aggregate-degree ranks (Supplementary Fig. 14, I, K, L), with *Operational SBM* visually closest to the observed pattern. *Independent SBM* had a smoother appearance (J). The bands arise when holdings share the same aggregate degree and hence average rank.

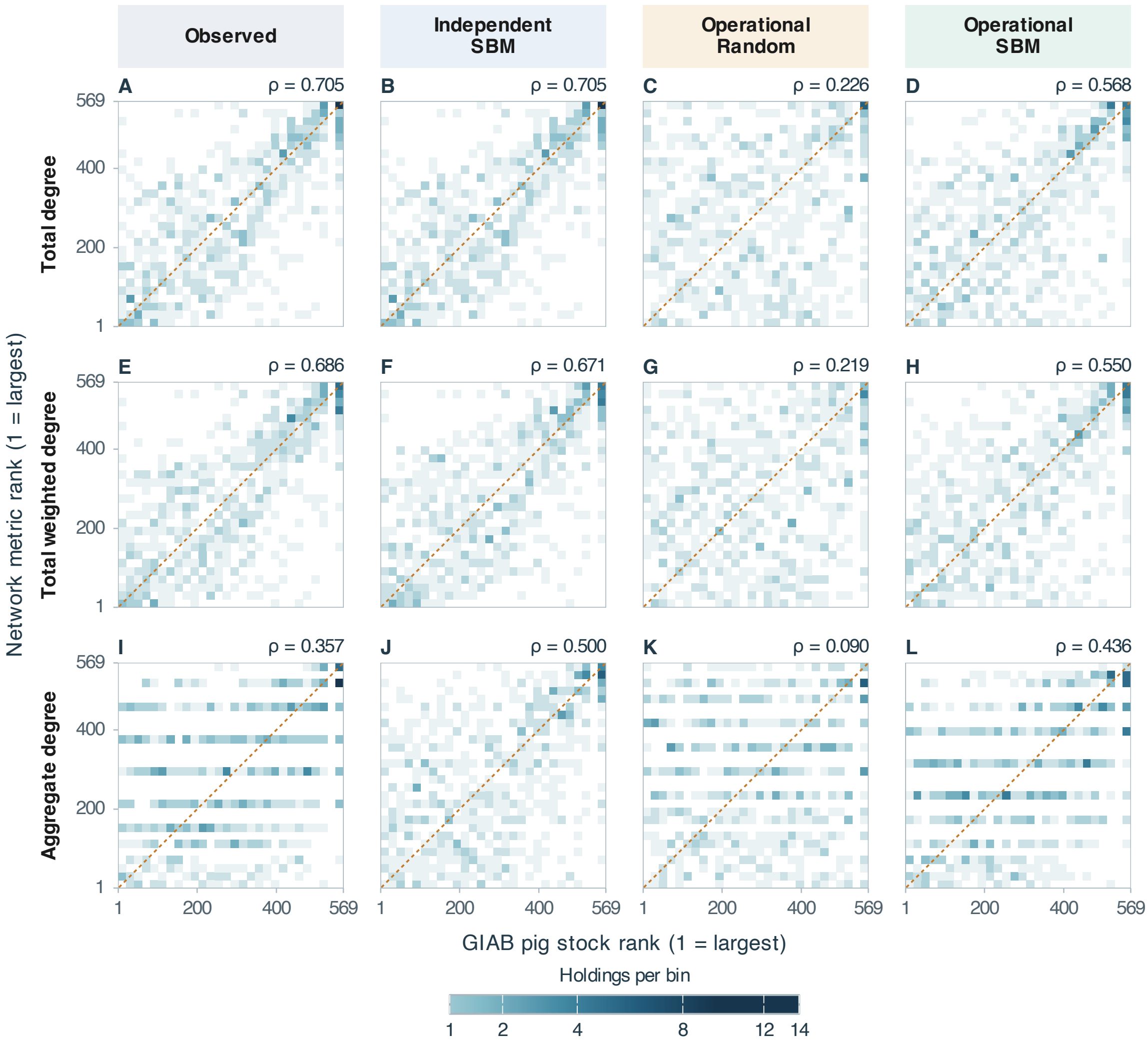


**Supplementary Fig. 14. Pig-stock ranks and holding-level activity across generation regimes.** The heatmaps compare GIAB pig-stock ranks with network-metric ranks for 569 holdings in the observed 2018 panel and the three synthetic versions. Panel rows show total degree (**A**–**D**), total weighted degree (**E**–**H**), and aggregate degree (**I**–**L**), respectively. Every panel uses the same $28 \times 28$ grid and square-root color scale for holdings per bin. Dashed diagonal lines mark equal ranks, and $\rho$ denotes Spearman correlation. Horizontal bands mainly arise from tied network-metric values. The aggregate-degree bands in the observed panel (**I**) reappear under both operational regimes (**K, L**), with *Operational SBM* visually closer to the observed pattern. Observed and *Independent SBM* total-degree panels coincide because holding degrees are preserved.

Larger tied groups can leave wider gaps between occupied ranks. This visual pattern provides a qualitative check of distinct trade partner organization alongside the stock–activity correlations and temporal reachability.

**Rank-based identity linkage.** We tested all three network activity metrics under two candidate populations. The first population contains 569 corresponding registry holdings and anonymous farm nodes to perform a known-membership benchmark. The second population contains all 633 CR35 registry holdings with positive pig totals and all 843 anonymous farm nodes to perform a full-roster benchmark. Between the GIAB registry and the anonymous network, there are 569 overlapping holdings. Furthermore, 64 GIAB holdings had no corresponding network node, and 274 network nodes had no recorded pig total in GIAB. The network nodes with no activity were still considered as candidates.

Rank-based record linkage has been used to assess re-identification risk in anonymized data [48]. Here, we used a benchmark that matches relative positions in pig-stock and network-activity rankings. Registry holdings and candidate nodes were sorted in descending order of pig stock and activity, respectively. For $M$ registry holdings and $N \geq M$ candidate nodes, we compared fractional positions $r/(M+1)$ and $j/(N+1)$, using the standard $k/(n+1)$ plotting positions [49]. Registry position $r$ was assigned to the nearest node position,

$$j(r) = \left\lfloor \frac{r(N+1)}{M+1} + 0.5 \right\rfloor, \qquad r = 1, \ldots, M,$$

which reduces to equal-position matching when $M = N$. Expected correct assignments were computed by averaging over independent uniform orderings within tied groups in both lists. Precision was the expected correct count divided by $M$. Under uniform random assignment, each of the $H$ shared identities had probability $1/N$ of a correct match, giving expected precision $H/(MN)$ (0.176% and 0.107% for the known-membership and full-roster benchmarks, respectively).

**Supplementary Table 5. Exact rank-linkage precision under two candidate populations.** Precision is the percentage of attempted registry-to-node assignments expected to be correct after averaging ties. Known membership uses 569 holdings and 569 nodes; the full roster uses 633 holdings and 843 nodes, with 569 corresponding identities. Degree counts daily contacts, weighted degree sums shipment sizes, and aggregate degree counts distinct partners separately by direction. Values are rounded to three decimal places.

| | Known membership | | | Full roster | | |
|---|---|---|---|---|---|---|
| Panel | Degree | Weighted | Aggregate | Degree | Weighted | Aggregate |
| Observed | 0.559% | 0.447% | 0.268% | 0.248% | 0.032% | 0.157% |
| *Independent SBM* | 0.559% | 0.204% | 0.286% | 0.248% | 0.190% | 0.124% |
| *Operational Random* | 0.271% | 0.048% | 0.231% | 0.232% | 0.029% | 0.150% |
| *Operational SBM* | 0.429% | 0.246% | 0.292% | 0.119% | 0.362% | 0.214% |
| Random assignment | 0.176% | 0.176% | 0.176% | 0.107% | 0.107% | 0.107% |

Overall, precision remained below 0.6% in every configuration (Supplementary Table 5). All synthetic panels matched or reduced total-degree precision in both populations and weighted-degree precision under known membership. In the full-roster experiment, weighted-degree precision increased from 0.032% observed to 0.190% for *Independent SBM* and 0.362% for *Operational SBM*. Aggregate-degree precision also increased in some of the comparisons.
